# A multigenic quantitative trait locus underlies natural variation in *Arabidopsis thaliana* root system architecture and transcriptional responses to microbiota-derived *Pseudomonas*

**DOI:** 10.1101/2025.09.03.673899

**Authors:** Charles Copeland, Elke Logemann, Milena Malisic, Anton Amrhein, Alessandro Valisi, Paul Schulze-Lefert

## Abstract

Plants interact with structured microbial communities called the microbiota, which can have a profound impact on plant growth and health. However, how plants perceive and respond to specific core microbiota members at a molecular level is still unclear. We identified natural variation in *Arabidopsis thaliana* root responses to bacterial strains of the genus *Pseudomonas*, a core genus of the plant microbiota. Some *A. thaliana* accessions such as Van-0 show strong root responses to *Pseudomonas* strains, including changes in root system architecture and transcriptional reprogramming. Through a forward genetic screen using *Pseudomonas* isolate R569, we found that the nuo NADH dehydrogenase complex, part of the bacterial electron transport chain, contributes to the bacterial activity on Van-0 roots. Using recombinant inbred lines, we further mapped a multigenic quantitative trait locus in the host that is associated with the root responses. In Van-0, the exocyst subunit *EXO70E2* positively contributes to the response, while Col-0 haplotypes of malectin-like and leucine-rich-repeat domain-containing receptor-like kinases play an inhibitory role. The identification of these components in the bacteria and the host establishes a genetic framework for how root developmental plasticity is integrated with the microbe-rich soil environment and is subject to intraspecific natural variation.

## Introduction

In the environment, plants interact with diverse microbial communities. Soil is especially microbe-rich, both in terms of microbial biomass and diversity, and plant roots thus come into contact with myriad microorganisms that perform important ecological functions (Banerjee and van der Heijden 2023). From these soil communities, plants impose a filtering step such that the microbial community closely associated with the root, termed the root microbiota, is less diverse than the surrounding soil biome, and is consistently dominated by similar taxa of microbes, termed core microbiota members (Copeland et al. 2025).

Plant roots perform a number of important functions, including uptake of soil-derived nutrients and water. Root system architecture (RSA) is highly plastic, responding to multi-factorial cues from the heterogenous soil environment. For example, limitation of some mineral nutrients leads to inhibition of primary root elongation (Kellermeier et al. 2014; Bouain et al. 2019), and lateral root emergence is promoted in soil patches with increased ammonium or water availability (Meier et al. 2020; Scharwies et al. 2025). RSA responses are subject to inter- and intraspecific natural variation. *Arabidopsis thaliana* natural accessions show differences in RSA, both under control conditions and in response to environmental variation (Lachowiec et al. 2015; Bouain et al. 2019; Deja-Muylle et al. 2022; LaRue et al. 2022).

The microbiota also exerts an important influence on RSA (Dini-Andreote et al. 2025). Many microbes can produce plant hormones or hormone mimics to influence plant growth. Additionally, microbes can also interact with plants through more complex mechanisms that interface with the plant’s endogenous signaling pathways. In many cases, the downstream phenotypic outcomes of these interactions, including improved plant stress tolerance, have been well characterized (Copeland et al. 2025). However, the upstream perception and signaling mechanisms remain to be identified.

Natural variation in plant responses to the microbiota can provide a means to identify the underlying genetic factors involved in plant-microbiota interactions. Microbiota-associated traits are frequently polygenic, with genome-wide association (GWA) studies identifying many associated polymorphisms, each with a small effect size, and low overlap between the significant polymorphisms for different traits (Wintermans et al. 2016; Roux et al. 2023). These GWA studies often consider plants within natural soil microbial communities, which increases complexity since both plant-microbe and microbe-microbe interactions occur in these systems. Given the complex genetic architecture underlying plant interactions with the microbiota, simplified experimental conditions, involving fewer plant genotypes grown in gnotobiotic systems with limited microbial diversity, may provide opportunities to identify and functionally validate important genes (Northen et al. 2024).

In this study, we investigated RSA changes in *A. thaliana* in response to microbiota-derived *Pseudomonas* strains, with natural genetic variation in both the plant host and the bacteria. Via an *in planta* forward genetic screen, we identified a bacterial gene that contributes to the root responses. We additionally cloned a host quantitative trait locus (QTL) containing multiple causal genes, which contribute both positively and negatively to the root responses.

## Results

To identify variation in plant responses to core microbiota members, we grew a panel of *Arabidopsis thaliana* natural accessions with *Pseudomonas* isolate R569, a member of the *At*-Sphere *A. thaliana* root culture collection (Bai et al. 2015). Previous studies have found that R569 suppresses root growth inhibition associated with pattern triggered immunity in *A. thaliana*, and has *in vitro* antagonistic activity against a broad range of microbiota members (Ma et al. 2021; Getzke et al. 2023). Given its multiple functions within the rhizosphere, we reasoned that R569 would be a good candidate to identify specific interactions with the plant host.

We observed variation in root system architecture (RSA) among the natural accessions grown in monoassociation with R569 (**Fig. 1a-g, Extended Data Fig. 1**). The reference accession Col-0 showed limited RSA changes in response to R569. However, many accessions showed some degree of inhibition in primary root growth, especially in the root zone where there are no lateral roots (the apical root), and increased lateral root density. These RSA changes were especially pronounced in a few accessions, such as Van-0. Despite the altered RSA, R569 does not affect the root or shoot biomass of Van-0, indicating that R569 influences root development in Van-0 without detectable growth-defense tradeoffs characteristic of infection by a pathogen (**Fig. 1h**). We therefore chose to focus on Col-0 and Van-0 as contrasting accessions to further characterize the R569-mediated root responses and identify their mechanistic basis.

**Fig. 1.**
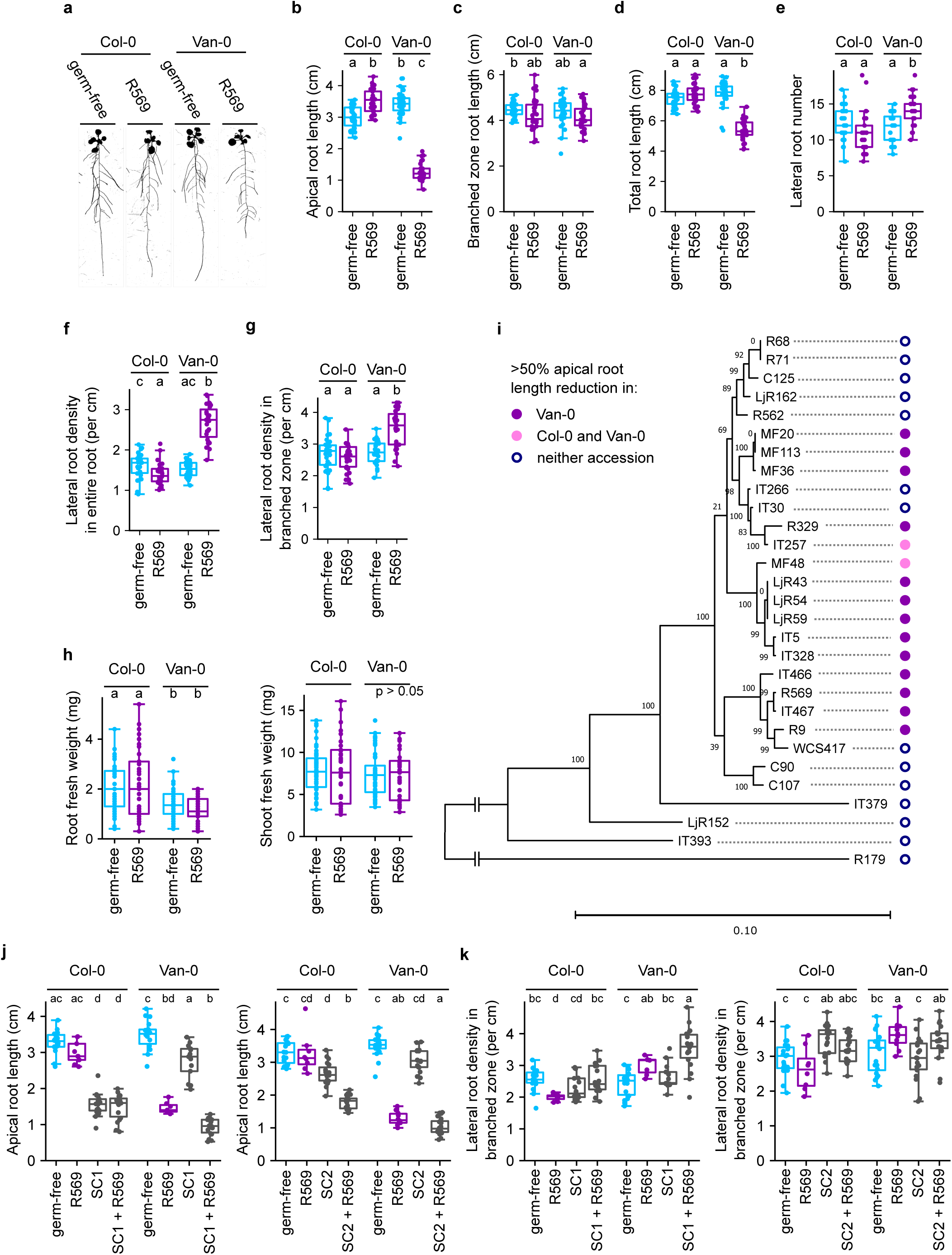
*A. thaliana* accessions Col-0 and Van-0 show differential root system architecture changes in response to root-associated *Pseudomonas* isolates. **a,** Binarized images of representative 16-day-old Col-0 and Van-0 plants grown in germ-free conditions or in monoassociation with *Pseudomonas* R569 for 7 days. **b-g,** Root architecture traits, showing apical root zone length **(b)**, branched root zone length **(c)**, total primary root length **(d)**, number of lateral roots **(e)**, lateral root density along the entire primary root **(f)**, and lateral root density in the branched zone **(g)** of accessions Col-0 and Van-0 grown in germ-free conditions or in monoassociation with *Pseudomonas* R569. **h,** Shoot and root fresh weight of 16-day-old plants of accessions Col-0 and Van-0 grown in germ-free conditions or in monoassociation with *Pseudomonas* R569 for 7 days. **i,** Phylogeny of microbiota-derived *Pseudomonas* isolates grown with Col-0 or Van-0, showing their effects on RSA. Phylogenetic relationships were inferred based on AMPHORA2 phylogenetic marker genes from the core genome. *Rhodanobacter* strain R179 was included as an outgroup. Strains were classified as causing strong RSA changes in Van-0 only (dark purple circles), both Col-0 and Van-0 (light pink circles), or neither accession (unfilled blue circles) based on the significant reduction of apical root length in each accession to less than 50% of the germ-free length (data shown in **Extended Data Fig. 2a**). **j, k,** Root architecture traits, showing apical root zone length **(j)** and lateral root density in the branched zone **(k)** of Col-0 and Van-0 grown in the presence of two independent bacterial synthetic communities, with or without R569. In the box plots, boxes represent the interquartile range (IQR) and whiskers represent 1.5 IQR beyond the boxes. The median is shown as a line within the box. Points represent root measurements from individual plants. Different letters represent significant differences (p < 0.05) according to a Kruskal-Wallis test followed by a Dunn’s posthoc test with a Benjamini-Hochburg False discovery rate correction for multiple testing.

To determine whether there is also natural variation in the ability of *Pseudomonas* isolates to influence RSA, we screened Col-0 and Van-0 with *Pseudomonas* isolates from different plant microbiota-derived culture collections (**Fig. 1i**, **Extended Data Fig. 2a**). The majority of tested isolates caused RSA changes in Van-0. Several isolates also triggered RSA responses in Col-0, consistent with previous reports (Finkel et al. 2020; Conway et al. 2022; Gonin et al. 2023). Variation in RSA responses was not explained by variation in the capacity of bacteria to colonize roots: despite their differential effects on root architecture, isolates R569 and WCS417 reached a similar bacterial load on roots of both Col-0 and Van-0 (**Extended Data Fig. 2b**). The accession-specific variation in RSA changes between Col-0 and Van-0 appears to be specific to a core clade of *Pseudomonas*, as we did not observe differential RSA responses when these accessions were grown with more taxonomically basal *Pseudomonas* strains, or with bacterial isolates representing diverse genera of the core root microbiota (**Fig. 1i**, **Extended Data Fig. 2a,c**).

To determine whether the *Pseudomonas*-mediated RSA changes are robust in a community context, we designed two independent synthetic communities (SynComs), each composed of 7 bacterial isolates representing core microbiota members. The isolates are part of the *At*-Sphere culture collection and thus originate from the same host plant species as R569. Van-0 roots showed similar RSA changes when simultaneously co-inoculated with R569 and each SynCom, compared to R569 alone (**Fig. 1j,k**). The accession-specific RSA changes that we observed in monoassociation are therefore retained when R569 is part of a bacterial community.

Microbiota-mediated inhibition of primary root growth has been attributed to microbial auxin biosynthesis (Finkel et al. 2020). However, Col-0 and Van-0 do not show differential RSA sensitivity to exogenous indole acetic acid (IAA) treatment (**Extended Data Fig. 3**), suggesting that R569 causes accession-specific RSA changes through a mechanism independent of bacteria-derived auxin. Additionally, neither heat-killed R569 nor cell-free supernatants of R569 induced root architecture changes in Van-0 (**Extended Data Fig. 4**), indicating that the accession-specific RSA changes require close association of live bacteria with the roots.

To determine whether the accession-specific RSA changes are maintained under varying growth conditions, we tested an additional gnotobiotic system based on perlite substrate (Ma et al. 2022). Consistent with our results on agar plates, R569 treatment also caused a reduction in root length in Van-0 but not Col-0 in this growth system (**Extended Data Fig. 5**).

In our assays, we did not observe severe alterations in root architecture in plants grown with *P. simiae* WCS417 (**Fig. 1i**), although this strain was previously reported to trigger RSA changes in Col-0 and other accessions (Zamioudis et al. 2013; Wintermans et al. 2016). However, when using the inoculation method described in these studies, which results in a high density of cells on the surface of the media, we could recapitulate the WCS417-mediated RSA changes in both accessions (**Extended Data Fig. 6**).

We next examined the temporal dynamics of the root growth response to R569, using a flood-inoculation method to avoid mechanical stress to seedlings that could affect root growth at early time points. Van-0 showed a significant inhibition in primary root growth rate starting at 48 hours post-inoculation (hpi), and the root growth rate remained low at later time points (**Fig. 2a**). Interestingly, Col-0 also showed a mild root growth inhibition at early time points, but root growth recovered at later time points. Neither accession showed a reduction in root growth following inoculation with WCS417.

**Fig. 2.**
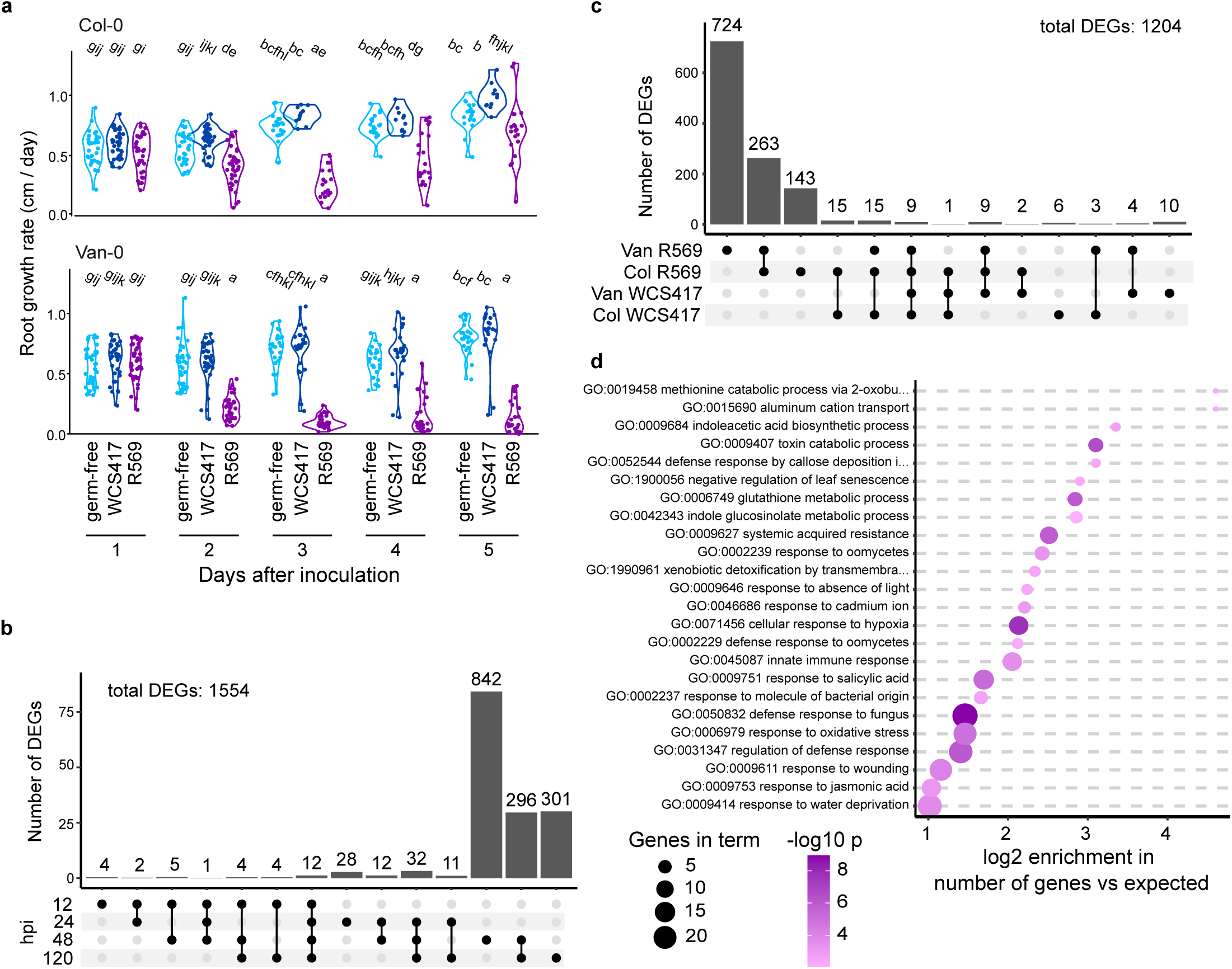
R569 triggers strong gene expression changes in Van-0 roots. **a,** Time course of primary root growth rate of seedlings flood-inoculated with the indicated bacterial strains. Plants were inoculated with bacterial strains by flooding 11 days after sowing. Col-0 and Van-0 were assayed together, but are shown on different axes for visual clarity. **b,** Upset plot showing number of DEGs at 12, 24, 48, and 120 hours post inoculation (hpi) with the indicated bacterial strains. Plants were inoculated at 11 days after germination. Genes with significant differential expression in at least one accession for at least one bacterial treatment compared to the germ-free control were counted as DEGs **c,** Upset plot showing the number of DEGs for each accession and bacterial treatment compared to germ-free control at 48 hpi. **d,** GO Biological Process terms enrichment in genes that show significantly stronger transcriptional upregulation by R569 treatment in Van-0 compared to Col-0 at 48 hpi. Significant DEGs were assessed based on the adjusted p < 0.05 for the interaction between genotype and bacterial treatment.

To further characterize the root responses to *Pseudomonas*, we examined the root transcriptome. We identified relatively few differentially expressed genes (DEGs) at the early 12 hpi and 24 hpi time points, with pronounced transcriptional changes first becoming apparent at 48 hpi (**Fig. 2b**, **Extended Data Fig. 7**). The highest number of DEGs was found in Van-0 treated with R569: at 48 hpi, 724 DEGs (60% of the total) were exclusive to this treatment combination, and another 22% of the DEGs showed differential expression in both accessions treated with R569 (**Fig. 2c, Supplementary Dataset 1a,b**). Consistent with the lack of RSA responses, WCS417 treatment caused relatively few gene expression changes, with only 10 significant DEGs unique to the Van-0 WCS417 treatment.

We performed a Gene Ontology (GO) enrichment analysis for genes whose expression was differentially affected by R569 treatment in Van-0 compared to Col-0 at 48hpi. Among genes showing relative upregulation in Van-0 treated with R569, we found an enrichment of GO terms related to innate immunity and defense (**Fig. 2d**). The general non-self response (GNSR) genes, a set of genes that are consistently upregulated by microbiota-derived bacteria in leaves (Maier et al. 2021), are significantly enriched in the set of upregulated DEGs: of the 24 GNSR genes, 15 are significantly induced by R569 treatment in Van-0 roots (p < 0.001), and 7 are induced by R569 in Col-0 roots (p < 0.001). In contrast, downregulated genes in Van-0 showed enrichment of GO terms related to root growth and cell wall biogenesis (**Extended Data Fig. 8**).

To identify bacterial factors involved in triggering RSA changes, we performed an *in planta* forward genetic screen using a *miniTn5* random insertional mutant library of R569 (Getzke et al. 2023). A strain carrying a *mTn5* insertion in the *nuoM* gene (R569 *nuoM::Tn5*) showed a marked reduction in its effect on Van-0 RSA (**Fig. 3a**, **Extended Data Fig. 9a**). An independent scarless deletion mutant of *nuoM* (R569 Δ*nuoM*) also shows reduced RSA-altering activity, confirming that loss of *nuoM* is causal for the reduction in RSA changes (**Fig. 3a**). The *nuo* operon encodes the NADH dehydrogenase complex I, a component of the bacterial electron transport chain that plays a major role in energy metabolism (Hreha et al. 2021; Ciemniecki and Newman 2023). The entire *nuo* operon, encoding subunits nuoA – nuoN, is well conserved in the genomes of microbiota-derived *Pseudomonas* isolates, and the sequence similarity does not correlate with the RSA changes triggered by the strains (**Extended Data Fig. 9b**). Although expression of *nuoM* is strongly reduced in both strains, the R569 *nuoM::Tn5* mutant also shows extremely low expression of *nuoN*, whereas expression in the scarless R569 Δ*nuoM* is not different from wild-type (**Extended Data Fig. 9c**). This polar effect of R569 *nuoM::mTn5* in the operon might explain why the phenotype of R569 Δ*nuoM* is not as strong.

**Fig. 3.**
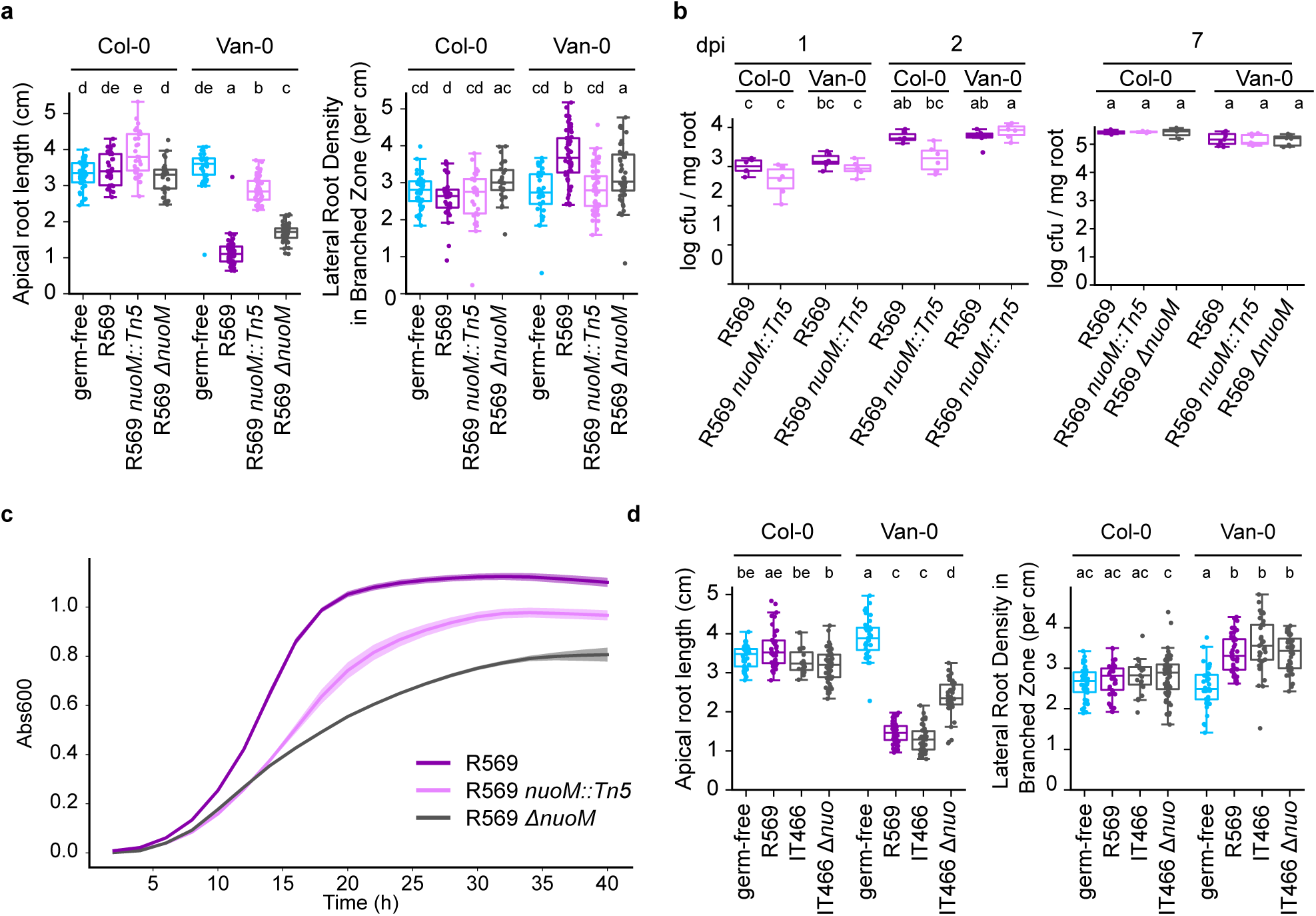
nuoM contributes to root architecture changes. **a,** Apical root length and lateral root density in the branched zone of Col-0 and Van-0 grown in germ-free conditions or in monoassociation with wild-type R569, R569 *nuoM::Tn5*, or R569 Δ*nuoM*. **b,** Bacterial load of R569, R569 *nuoM::mTn5*, or R569 Δ*nuoM* on roots of Col-0 or Van-0 at 1, 2, and 7 days post inoculation (dpi). **c,** Growth curves of R569, R569 *nuoM::mTn5*, or R569 Δ*nuoM* grown in liquid culture in rich media. Lighter colored bands around the lines indicate 95% confidence intervals. **d,** Apical root length and lateral root density in the branched zone of Col-0 and Van-0 grown in germ-free conditions or in monoassociation with R569, wild-type IT466, or a IT466 Δ*nuo* scarless operon deletion mutant.

*Pseudomonas. aeruginosa* PAO1 is known to produce two additional NADH dehydrogenases, nhd and the nqr complex, which perform the same biochemical function as the nuo complex (Hreha et al. 2021; Ciemniecki and Newman 2023). Of these, *nhd* was also present in the genomes of all tested *Pseudomonas* isolates, while a full *nqr* operon was only found in one isolate (**Extended Data Fig. 9b**). R569 *nuoM* mutants do not show defects in colonization on Van-0 roots, indicating that *nuoM* is not required for R569 to grow and colonize the root compartment in mono-association (**Fig. 3b**).

However, *nuoM* mutants do have a slightly slower growth rate in rich media and reach a lower cell density at the stationary phase, indicating that loss of the *nuo* operon leads to metabolic defects under specific growth conditions (**Fig. 3c**). To establish whether the *nuo* operon is involved in accession-specific RSA changes caused by other *Pseudomonas* isolates, we generated a scarless deletion of the *nuo* operon in *Pseudomonas* isolate IT466 (IT466 Δ*nuo*). IT466 Δ*nuo* caused less severe root growth inhibition on Van-0 than the wild-type IT466 (**Fig. 3d**).

To identify the plant genetic factors responsible for the accession-specific root responses to R569, we crossed Col-0 with Van-0 and examined the root phenotypes of the offspring. F2 progeny from this cross showed a broad distribution in apical root length when grown with R569, and most plants had a phenotype intermediate between the two parent accessions (**Extended Data Fig. 10a**). A similar pattern was observed when Col-0 was crossed with Vim-0 and Ees-0, two other accessions that closely phenocopy Van-0 (**Extended Data Fig. 10b**). Based on the lack of a Mendelian 3:1 phenotypic segregation pattern, we concluded that the variation in RSA changes likely involves multiple loci that may show incomplete dominance. The variation in F2 phenotype hindered our ability to confidently score RSA traits.

Offspring from Van-0 x Vim-0 and Van-0 x Ees-0 crosses showed consistent RSA changes similar to the parental accessions, suggesting that the root responses in these three accessions involve allelic polymorphisms (**Extended Data Fig. 10b**).

To account for the apparently complex genetic architecture of this trait, we performed QTL mapping using a population of recombinant inbred lines (RILs) generated between Col-0 and Van-0 (Gerald et al. 2014). We identified a highly significant QTL on chromosome 5 that was associated with the accession-specific R569-mediated primary root growth inhibition (**Fig. 4a**). The QTL as a whole had a large effect size: the mean relative root length of RILs with the Van-0 haplotype at markers CoVa25 and CoVa32 was 1.36 standard deviations lower than for lines with the Col-0 haplotype at both markers (Cohen’s d = 1.36, **Extended Data Fig. 11a**). However, RILs with recombination points within the QTL region tended to show mild RSA changes in response to R569, more similar to Col-0 (**Extended Data Fig. 11**), regardless of whether the Van-0 haplotype was found at the telomeric or centromeric part of the QTL. This suggested that the QTL region may contain multiple causal genes, and the strong effect size of the QTL requires the Van-0 haplotype at all of them.

**Fig. 4.**
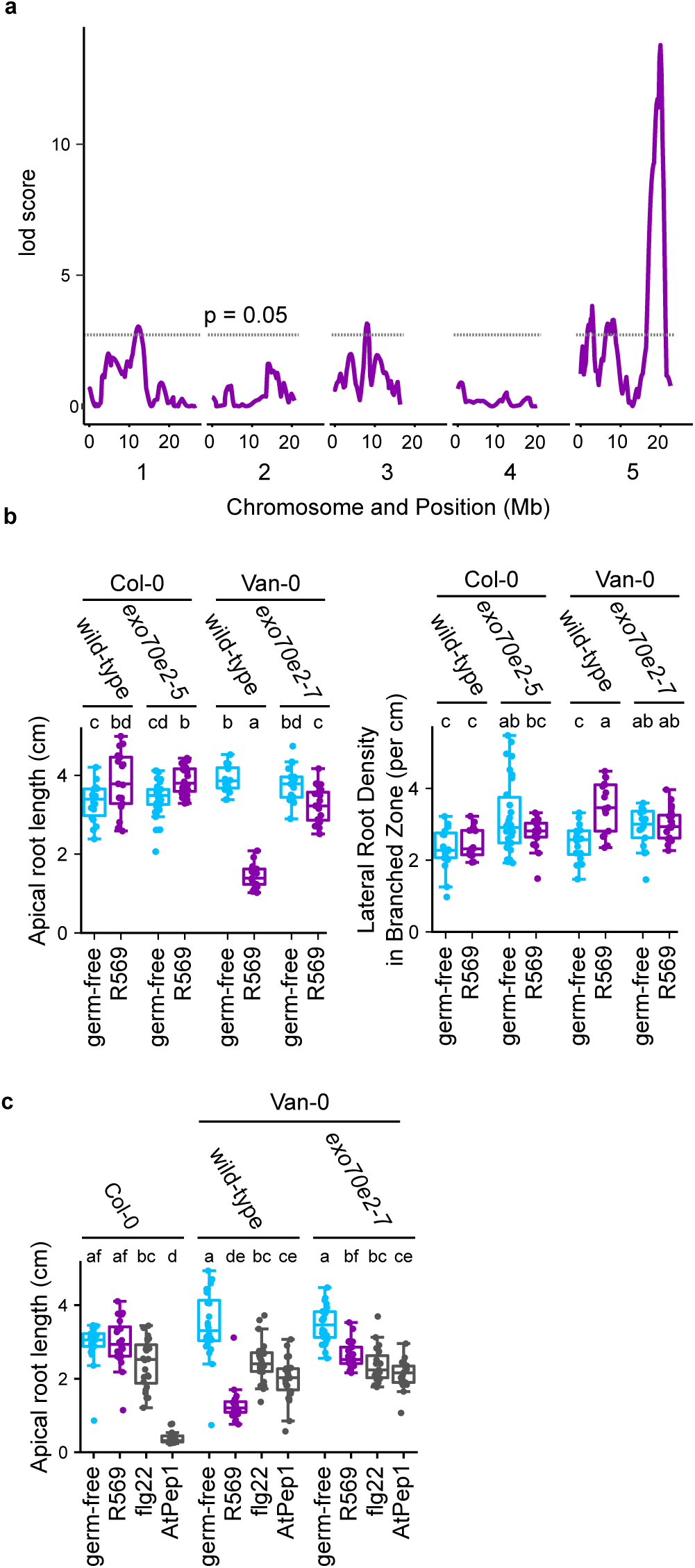
R569-triggered root architecture changes are associated with a quantitative trait locus on chromosome 5 that includes *EXO70E2*. **a**, lod scores from QTL mapping based on relative root length of a RIL population. The dashed grey line represents a significance threshold of p = 0.05. **b,** Apical root zone length and lateral root density in the branched zone of *exo70e2* mutants in the Col-0 and Van-0 background, grown in germ-free conditions or in monoassociation with R569. **c,** Apical root zone length of the *exo70e2-7* mutant grown with elicitors flg22 or AtPep1, or with R569.

To identify specific causal loci, we chose candidate genes within the QTL that have sequence polymorphisms between Col-0 and Van-0 and that showed accession-specific gene expression patterns or transcriptional responses to R569 treatment. We targeted these candidate genes using crispr-Cas9, and identified lines with genetic lesions in *EXO70E2* (*AT5G61010*) in the Van-0 background that do not show the dramatic R569-mediated RSA changes observed in the wild-type Van-0 parents (**Fig. 4b**).

*EXO70E2* is required for callose deposition in response to the bacterial elicitor flg22 in leaves (Redditt et al. 2019). To determine whether *EXO70E2* is also involved in pattern-triggered immune responses in roots, we grew *exo70e2* mutants with flg22 or the damage-associated molecular pattern AtPep1, which both cause primary root growth inhibition (RGI) (Poncini et al. 2017). In response to the elicitors, *exo70e2* mutants showed a similar degree of RGI compared to their respective wild-type parental accessions (**Fig. 4c**, **Extended Data Fig. 12a**). Therefore, *EXO70E2* is required for accession-specific RSA changes mediated by *Pseudomonas* strains such as R569, but not immunity-associated changes in root growth that occur in both accessions. Similarly, *EXO70E2* is not required for RSA changes triggered by *Pseudomonas* isolates that affect both accessions, or by exogenous IAA treatment, again supporting a role for *EXO70E2* in mediating accession-specific root responses, but not responses that occur in both accessions (**Extended Data Fig. 12b**)

As bacterial *nuoM* mutants and Van-0 *exo70e2* mutants are associated with weaker RSA changes, we performed an additional RNA-seq experiment to examine the impact of these mutations on R569-mediated transcriptional changes. Consistent with our previous results, Van-0 showed strong transcriptional reprogramming when grown with wild-type R569 compared to germ-free conditions (**Fig. 5a,b, Supplementary Dataset 1c,d**). The R569 *nuoM::Tn5* resulted in weaker transcriptional changes than the wild-type bacteria, with only 38% of the R569-regulated DEGs also regulated by the mutant bacteria. Loss of *EXO70E2* had a greater impact on plant responses to bacteria: the *exo70e2-7* mutant grown with wild-type R569 showed 15% of the DEGs identified in wild-type Van-0, and of these, 50% showed significantly weaker induction. The transcriptional responses were further suppressed by a combination of the two mutants, with Van-0 *exo70e2-7* plus R569 *nuoM::Tn5* showing only weak transcriptional changes. The two mutations showed a significant synergistic interaction for 21% of the R569-upregulated genes, where the expression was reduced more strongly in *exo70e2-7* treated with R569 *nuoM::Tn5* than can be explained by the additive effects of each mutation alone (**Supplementary Dataset 1e**). Thus, bacterial *nuoM* and *EXO70E2* in the host both contribute to transcriptional reprogramming of partially overlapping sets of host genes.

**Fig. 5.**
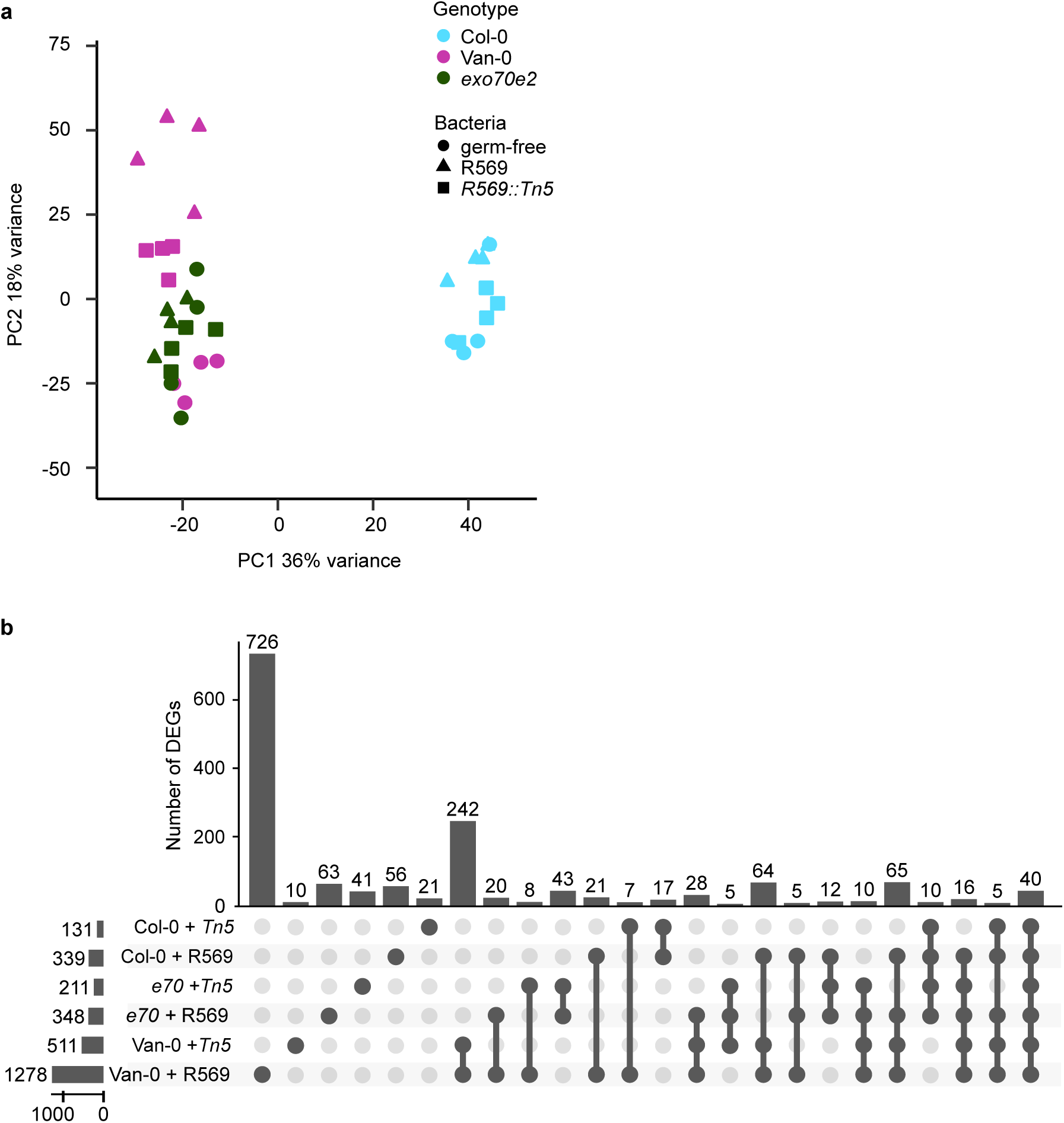
*nuoM* and *EXO70E2* both contribute to root transcriptional responses to R569. **a,** PCA plot of root transcriptomes of 11-day-old Col-0, Van-0, and Van-0 *exo70e2-7* grown in germ-free conditions or in monoassociation with R569 or R569 *nuoM::Tn5* for 48 hours. **b,** Upset plots showing numbers of DEGs in the roots of 11-day-old plants of the given genotypes grown with R569 or R569 *nuoM::Tn5* for 48 hours, relative to each genotype grown in germ-free conditions. For clarity of visualization, only sets with at least 5 DEGs are shown. *e70* refers to *exo70e2-7,* T*n5* refers to R569 *nuoM::Tn5*.

Of the 24 GNSR genes, 18 were significantly induced in Van-0 with wild-type R569, and the majority of these were also significantly upregulated in the Van-0 plus R569 *nuoM::Tn5* or *exo70e2-7* plus R569 treatment combinations (15 and 13 genes, respectively), albeit to lower levels. By contrast, only one GNSR gene was significantly upregulated in the *exo70e2-7* plus R569 *nuoM::Tn5* condition.

Given the genetic evidence that the QTL may contain multiple causal loci, we sought to identify additional genes regulating the differential root responses. Plant responses to microbes are often mediated by pattern recognition receptors on the cell surface. Within the QTL region, we identified a tandem cluster of four genes encoding Receptor-like Kinases (RLKs) with extracellular malectin-like (MLD) and leucine-rich repeat (LRR) domains, corresponding to Col-0 genes *AT5G59650 – AT5G59680*. Natural structural variation in this cluster has been described previously, as a duplication of *AT5G59670* in some *A. thaliana* accessions has generated the hybrid incompatibility gene *OAK* (Smith et al. 2011). The Col-0 and Van-0 haplotypes of this gene cluster are syntenic with each other, and both lack the duplication in *AT5G59670* that led to *OAK*. For consistency in the literature, we therefore here refer to *AT5G59650 – AT5G59680* as *OAK-LIKE1* (*OAKL*) - *OAKL4* (**Extended Data Fig. 13a**). Despite retained gene synteny, the *OAKL* gene cluster is highly sequence-divergent between Col-0 and Van-0, including in the gene bodies and promoter regions (**Extended Data Fig. 13a,b**). Closely related MLD-LRR-RLK genes of the *Stress-Induced Factor* (*SIF*) family do not show such an extreme degree of sequence divergence between Col-0 and Van-0 (**Extended Data Fig. 13a**). Further, *OAKL* genes show accession-specific expression patterns in roots: *OAKL1* shows high expression in Van-0 roots and low expression in Col-0 roots, while *OAKL4* shows higher expression in Col-0 roots (**Extended Data Fig. 13c**). *OAKL4* is also upregulated in response to R569. The other MLD-LRR-RLK genes show varying levels of constitutive expression in roots, but their expression is similar between accessions and not significantly affected by the bacterial treatments. We thus selected the *OAKL* cluster as a candidate to test with CRISPR-Cas9 gene editing.

Gene-editing of *OAKL* genes in the Col-0 and Van-0 background, including large deletions inactivating all four *OAKL* genes (*oakl-clus*), did not alter the root architecture responses to R569 compared to their respective wild-type parents (**Fig. 6a,b**, **Extended Data Fig. 14**). Thus, the Van-0 haplotypes of *OAKL* genes likely do not contribute positively to the R569-mediated root responses in Van-0.

**Fig. 6.**
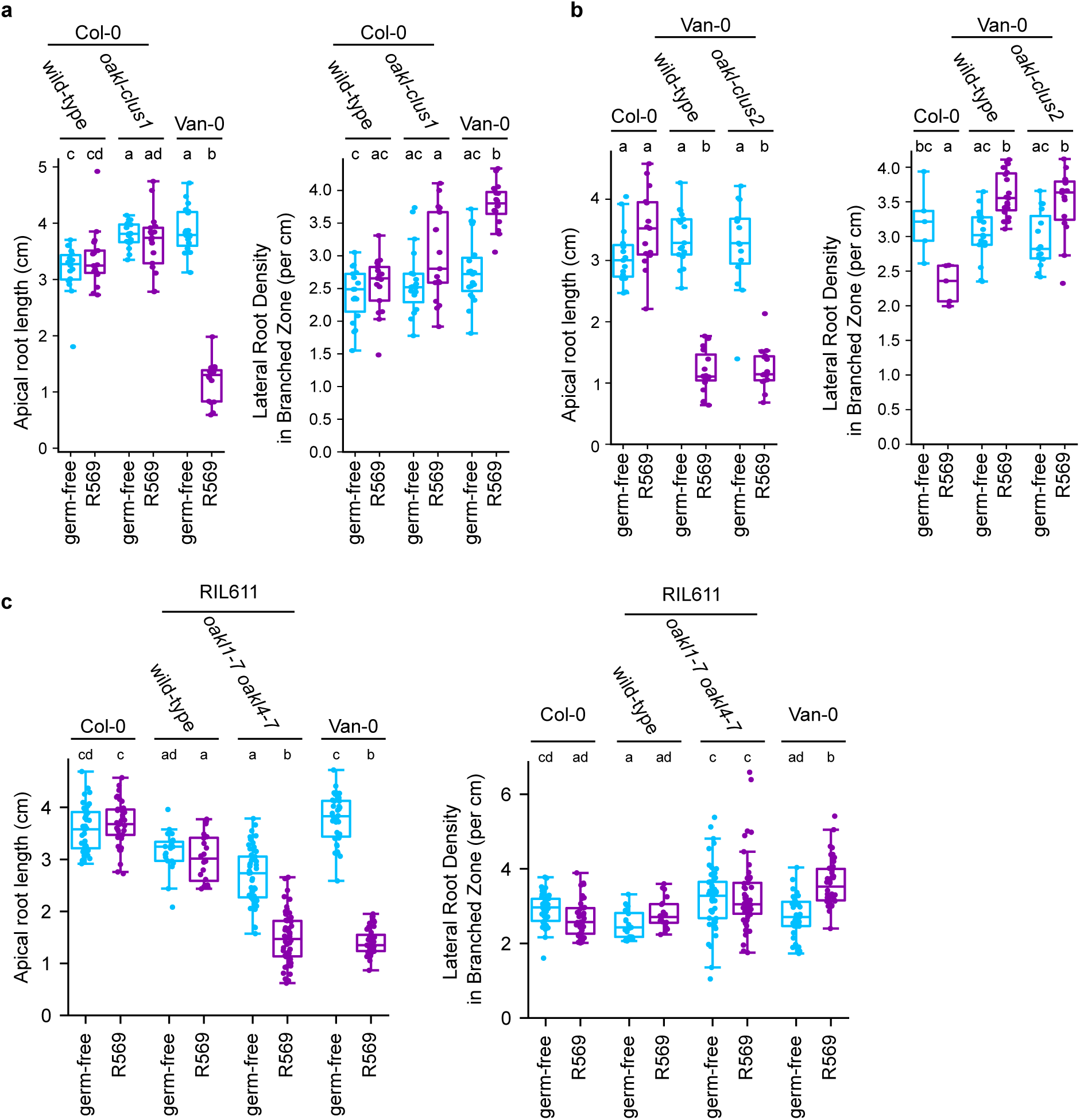
*OAKL* malectin-like domain and leucine-rich-repeat containing RLKs suppress R569-mediated root architecture changes. **a-b**, Apical root zone length and lateral root density in the branched zone of gene-edited lines with deletion of the *OAKL* gene cluster in the Col-0 **(a)** or Van-0 **(b)** background, grown in germ-free conditions or in monassociation with R569. **c,** Apical root zone length and lateral root density in the branched zone of a gene-edited line with edits in *OAKL1* and *OAK4* in the RIL611 background.

The dynamics of the root growth rate in Col-0 treated with R569, where a transient slowing of root growth is followed by a recovery to normal growth rate (**Fig. 2a**), suggested that Col-0 may have an active mechanism that suppresses the root responses in this accession. We reasoned that knocking out candidate genes through CRISPR-Cas9 gene editing in Col-0 background may not be sufficient to identify genes involved in this mechanism, since the Col-0 background lacks the additional factors in the QTL, such as the Van-0 haplotype of *EXO70E2*, which are required to trigger strong root responses.

To determine whether the *OAKL* cluster is involved in negative regulation of root responses to R569, we therefore edited the *OAKL* genes in the genetic background of RIL611, which has had a recombination event within the QTL region resulting in the Col-0 haplotype in the centromeric part of the QTL, including the *OAKL* cluster, and the Van-0 haplotype in the telomeric region of the QTL that includes *EXO70E2* (**Extended Data Fig. 11b**). Although the wild-type RIL611 parent shows relatively mild root responses to R569, a line with gene edits in *OAKL1* and *OAKL4* shows a strong R569-mediated reduction in apical root length (**Fig. 6c**), supporting an inhibitory activity. Editing of *OAKL1* alone is not sufficient to confer strong root responses in the RIL611 background (**Extended Data Fig. 14**). We found the Col-0 haplotype of *OAKL4* to be recalcitrant to gene editing compared to *OAKL1*, or the Van-0 *OAKL4* haplotype (**Supplementary Table 1**), and have not yet been able to assess the impact of editing only *OAKL4* but not *OAKL1* in the RIL611 background.

## Discussion

The electron transport chains of *Pseudomonas* are highly branched, with the genome of *P. aeruginosa* encoding 3 NADH dehydrogenase enzymes (Hreha et al. 2021). Mutants with deletions of any one or even two of these enzymes often grow well in rich media. This is consistent with our observation that R569 *nuoM* mutants do not show defects in root colonization. However, the importance of each NADH dehydrogenase varies depending on availability of oxygen and different carbon sources (Torres et al. 2019). The reduced stationary-phase cell density of R569 *nuoM* mutants in rich media suggests that these mutants may have a reduced ability to use specific metabolites. We hypothesize that the nuo NADH dehydrogenase complex is involved in specific metabolic activities of R569 *in planta* that are required for triggering strong root responses in a subset of Arabidopsis accessions, but are dispensable for robust bacterial growth in the root compartment.

Using a population of advanced recombinant inbred lines, we have cloned a host QTL in which natural variation at multiple linked genes leads to variation in plant responses to a core microbiota member.

One of the genes we identified in the QTL region is *EXO70E2*, which is required for the RSA changes and transcriptional responses to R569 in Van-0. EXO70 is a subunit of the exocyst complex, a conserved complex involved in eukaryotic protein secretion (De la Concepcion 2023). In plants, the *EXO70* gene family has diversified, and several family members are involved in plant-microbe interactions, often by specifically affecting the abundance or distribution of plasma membrane-localized effectors or controlling cell wall deposition (De la Concepcion 2023). In leaves, EXO70H4 interacts with MLO proteins, including the GNSR MLO12, to mediate targeted deposition of callose and cell wall carbohydrate in trichomes, and are also required for susceptibility to powdery mildew (Huebbers et al. 2024). *EXO70B1* and *EXO70B2* also influence transcriptional responses to leaf microbiota (Keppler et al. 2025). The *A. thaliana exo70b1 exo70b2* double mutant shows distinct transcriptional regulation in response to two taxonomically unrelated leaf commensals that trigger overlapping responses in wild-type Col-0, indicating that these subunits contribute to convergence of host responses upon detection of specific microbial strains.

Canonically, EXO70 subunits are thought to bind phospholipids at the plasma membrane, linking secretory vesicles to the plasma membrane before fusion, thereby remodeling the composition of membrane proteins on the cell surface and extracellular cargo in the apoplast (Žárský 2022).

However, several EXO70 family members have been shown to have non-canonical functions that do not require the full exocyst complex. EXO70E2 has been found in Exocyst-Positive Organelles (EXPOs), double membrane-bound structures that are involved in unconventional protein secretion (Wang et al. 2010; Ding et al. 2014). Although cargo selectivity of EXPO-mediated secretion is unclear, cell wall-related proteins have been localized to EXPOs, suggesting that EXO70E2-mediated secretion may be important for modifying cell wall properties (Wang et al. 2010; Poulsen et al. 2014). *EXO70E2* was recently reported to be important for production of specific populations of extracellular vesicles (Koch et al. 2025). Thus, while Van-0 *EXO70E2* likely facilitates responses to and/or perception of *Pseudomonas* through targeted secretion, it remains unknown which cargoes are involved, and whether they are trafficked to the cell surface and/or secreted into the apoplast.

Another component of the QTL identified here is the *OAKL* cluster of genes encoding MLD-LRR-RLKs. Col-0 and Van-0 share a genomic architecture at this locus, although the genes themselves are highly polymorphic. However, the locus has diversified even further in other accessions through gene deletions and duplications (Smith et al. 2011). Indeed, the originally described *OAK* gene in accessions Sha and Bla-1 likely arose through a duplication of *OAKL3*, and is not present in either Col-0 or Van-0; meanwhile, *OAKL1* has been lost in Bla-1. The diversification of MLD-LRR-RLKs may be related to their heterodimer formation. *OAK* was identified through a hybrid incompatibility phenotype observed when the Sha and Bla-1 haplotypes are brought into the same plant through crossing or transgenic expression (Smith et al. 2011). Heterodimerization was also reported in a tandem MLD-LRR-RLK pair from *Medicago*, and accessions containing functional copies of both genes failed to form symbiotic relationships with *Sinorhizobium* bacteria (Liu et al. 2022).

Loss of the *OAKL* function did not lead to R569-mediated RSA changes in the Col-0 background, but only in a genetic background containing the Van-0 haplotype in the telomeric region of the QTL, which includes *EXO70E2*. A lineage-specific *EXO70FX12* family member was also identified as part of a genetic module, which defines the *RPS8* locus conferring barley stripe rust resistance, with the other gene being an RLK lacking a malectin-like domain (Holden et al. 2022). This module at *RPS8* generally shows a presence/absence pattern of polymorphisms among barley accessions, and *EXO70FX12* and the RLK both positively contribute to resistance, in contrast to our results where *EXO70E2* and the *OAKL* genes show opposite contributions.

While we do not know the identity of the ligands for the *OAKL* genes, MLD-containing RLK proteins commonly bind to components of the plant cell wall and are involved in plant immune signaling (Yang et al. 2021). The LRR-MD-RLKs CORK1, IGP3, and IGP4 bind to cellulose or xylan oligomers, and are required for immune activation upon treatment with these DAMPs (Tseng et al. 2022; Martín-Dacal et al. 2023; Fernández-Calvo et al. 2024). FERONIA interacts with de-esterified pectin and RALF peptides, and was recently shown to promote recycling of membrane proteins under stress conditions that threaten cell wall integrity via endocytosis (Malivert and Hamant 2023; Liu et al. 2024). Among the reported phenotypes of *fer* mutants is increased abundance of *Pseudomonas* in the rhizosphere (Song et al. 2021). The LRR extensin proteins LRX1 and LRX2, which interact with FER and the cell wall (Ortiz-Morea et al. 2021), similarly negatively regulate the relative abundance of rhizosphere *Pseudomonas* (Song et al. 2025). Although their ligands remain unknown, MLD-LRR-RLKs SIF2 and IOS1 also interact with known immune signaling components and contribute to disease resistance (Yeh et al. 2016; Yuan et al. 2018; Chan et al. 2020). It is thus tempting to speculate that plants may indirectly perceive microbiota members partly through their effects on cell wall integrity, and that natural variation at the *OAKL* locus may lead to differences among natural accessions for their tolerance to cell wall integrity perturbations. Given the high degree of intra-species sequence divergence in the *OAKL* gene cluster, including presence/absence polymorphisms, we hypothesize that *OAKLs* are unlikely to perform ‘hard-wired’ developmental roles, but rather serve to integrate root developmental processes with cues from the microbe-rich soil environment along with *EXO70E2*.

Our study has shown the power of advanced RILs to dissect the complex genetic architecture underlying plant interactions with the microbiota under simplified experimental conditions. Recent advances in the understanding and genetic control of meiotic recombination in plants has made it possible to massively increase meiotic recombination frequencies in *A. thaliana*, accelerating the generation of RILs for the detection of gene-scale QTLs with only a few hundred plants (Capilla-Pérez et al. 2024). This can expedite the identification of plant factors underlying the complex genetics of plant-microbiota interactions.

## Methods

### Plant material and growth conditions

*Arabidopsis thaliana* seeds were briefly surface sterilized in a 2% sodium hypochlorite solution, washed thoroughly with sterile water, and resuspended in 0.1% agar for planting. Except where noted, seeds were pregerminated on 120 cm square plates (Grenier) filled with ½ strength Murashige and Skoog (MS) media including vitamins (Duchefa Biochemie, Haarlem, NL) with 0.5% sucrose, solidified with 1.1% Bacto agar (Catalog #214010, BD). After 9 days, seedlings were transferred to new ½ MS plates solidified with a mixture of Bacto agar and Gelrite (Duchefa) at a 3:1 ratio, which increases the gelling strength and transparency of the matrix. Plates were kept vertically in a light cabinet (Panasonic) with a cycle of 10h light at 21°C / 14 h dark at 19°C.

Bacterial isolates were grown in ½ strength TSB media, washed, resuspended in 10 mM MgCl_2_, and added to the media at a final OD_600_ of 5 x 10^-4^ just before pouring plates. An equivalent amount of sterile MgCl_2_ was added for the germ-free controls. IAA and elicitor treatments were similarly added to the media before pouring. The final concentration of flg22 was 500 nM the final concentration of AtPep1 was 50 nM. For heat-killed bacteria, the bacterial suspension was incubated at 99°C for 10 minutes before adding to the growth media to a final OD_600_ of 1 x 10^-3^.

For SynCom experiments, individual isolates in the SynCom (**Supplementary Table 2**) were prepared as above and combined in equal cell densities, and the SynCom was added to the growth media to a total SynCom bacterial OD_600_ = 5×10^-4^. For simultaneous co-inoculation, R569 was additionally added to OD_600_ = 5×10^-4^.

Plates were scanned and lateral roots were counted from the scanned images. The primary root length, including length of the apical and branched root zones, was measured using the Fiji distribution of ImageJ (Schindelin et al. 2012).

### Phylogeny of *Pseudomonas* isolates

The *Pseudomonas* isolates were derived from multiple culture collections which is indicated by the strain prefix (**Supplementary Table 3**) (Lamers et al. 1988; Bai et al. 2015; Levy et al. 2018; Wippel et al. 2021; Durán et al. 2022). AMPHORA2 was used to extract the sequence of core phylogenomic marker genes from assembled genomes of culture collection isolates and to align the deduced amino acid sequences (Wu and Scott 2012). The maximum-likelihood tree was generated using FastTree (Price et al. 2009).

### Colony count assays

Roots were washed for 40 s and 10 s in sterile MgCl_2_, dried, and homogenized in MgCl_2_ using Precellys Evolution homogenizer (Bertin). Serial dilutions were plated on ½ TSB plates, and colonies were counted once they became visible.

### Cell-free supernatants

Seeds were sown on ½ MS plates without sucrose, solidified with Bacteriological agar (Catalog #214530, BD). After 11 days, plates were flooded with a bacterial suspension in ½ MS at an OD_600_ of 5 x 10^-4^, with sterile MgCl2 for the germ-free control. After 6 days incubation in the light cabinet, free liquid on the plates was collected, centrifuged briefly to remove most of the bacterial cells, and filter sterilized using a 0.22 μM filter (Millipore). The resulting cell-free supernatants were mixed with 2x volume of fresh ½ MS solidified with Bacteriological agar. Before filtering, the bacterial cultures had an OD_600_ of approximately 0.12; thus, if the cells had not been removed, the new plates would be approximately OD_600_ = 0.04. Seedlings that had been pregerminated as usual for 9 days were transferred to the new plates, and root architecture was analyzed 7 days after transfer.

### Perlite growth system

Experiments with the perlite gnotobiotic growth system were performed according to (Ma et al. 2022). Briefly, washed perlite was placed in round plates and 25mL of ½ MS medium, with vitamins, containing the bacterial suspension at OD_600_ = 5×10^-4^ or MgCl_2_ for germ-free treatment was added.

Seeds were planted on the surface of the perlite, and a second plate bottom was used as a lid. After 3 weeks of growth, plants were removed from the perlite by addition of excess water, roots were carefully cleaned and root length was measured.

### Van-0 genome *de novo* assembly

Van-0 high molecular weight genomic DNA was isolated using a DNeasy Plant Minikit (Qiagen). Sequencing was performed by the Max Planck-Genome-centre (Cologne, Germany https://mpgc.mpipz.mpg.de/home/) using the PacBio Sequel II platform. HiFi reads were assembled using Flye version 2.9 (Kolmogorov et al. 2019).

For comparison of the *OAKL* cluster, the genomic regions from Col-0 and Van-0 were aligned with nucmer program in MUMmer4 (Marçais et al. 2018).

### RNA-seq

For RNA-seq presented in **Fig. 2**, seeds were sown on ½ MS plates without sucrose, solidified with Bacteriological agar. After 11 days, plates were gently flooded with bacteria suspended in liquid ½ MS at OD_600_ of 5 x 10^-4^, and the excess liquid was removed after approximately 1 minute, and the plates were returned to the light cabinet. At the indicated time points after inoculation, whole roots were collected, washed briefly in water, and flash frozen in liquid nitrogen. Roots of approximately 40 seedlings were pooled for each biological replicate. The inoculations and harvesting were performed 6 hours after lights on in the growth cabinet, except for the 12 hpi time point. Total RNA was extracted using an RNeasy Plant Mini Kit (Qiagen) including DNase I treatment (Qiagen).

For the RNA-seq shown in **Fig. 5**, plants were grown and inoculated by seedling transfer as usual. Whole roots were collected at 48 hpi, washed briefly in water, and flash frozen in liquid nitrogen. Roots of approximately 40 seedlings were pooled for each biological replicate. Total RNA was extracted using TRIzol Reagent (Catalog # 15596018, Thermo) and DNA was removed with a DNA-free DNA Removal Kit (Catalog #AM1906 Thermo).

For both RNA-seq experiments, mRNA library preparation and sequencing were performed on an Illumina Novoseq platform by Novogene (Cambridge, UK).

Reads were trimmed with fastp (Chen 2023), using a quality threshold of 25 and a window length of 5. Reads were then aligned to the Col-0 reference genome (TAIR10) with the HISAT2 (Kim et al. 2019) using default settings, and counts were assigned using featureCounts (Liao et al. 2014). Genes within the QTL that are highly polymorphic between Col-0 and Van-0 were manually annotated in the Van-0 genome, and alignments of Van-0 reads to the Van-0 genome were used to assign counts for these genes. Differentially gene expression was analyzed with DESeq2 (Love et al. 2014).

GO enrichment analysis was performed using TopGO (Alexa and Rahnenfuhrer 2025).

### Bacterial forward genetic screen

A *minTn5* random insertion mutant library of R569 (Getzke et al. 2023) was screened in a phytostrip gnotobiotic system described in (Ma et al. 2022). Briefly, phytostrips were filled with ½ MS with vitamins, solidified with phytagel, with bacterial mutants suspended at OD_600_ = 5×10^-4^. Seeds were planted on the surface of the media, and the phytostrips were placed in a sealed microbox and incubated in the light cabinet.

### Generation of scarless R569 Δ*nuoM* mutant

The R569 Δ*nuoM* mutant was generated using the Golden Gate-compatible pOGG2 vector and primers L979, L980, L981 and L982 in a first step as described previously (**Supplementary Table 4**) (Ordon et al. 2025).

To increase the sucrose counter selection efficiency the assembled flanking regions were cloned into the pEXG2 vector (Melnyk et al. 2019). In brief, the assembled flanking regions from pOGG2 *nuoM* were PCR amplified with primers L983 amd L984 including BamHI and HindIII restriction sites and ligated into the BamHI and HindIII sites of pEXG2. Ligation was transformed into chemically competent aliquots of the diaminopimelic acid (DAP) auxotroph *E. coli* BW29427 and plated on LB containing 0.3mM DAP and 50 μg/mL gentamycin.

To conjugate pEXG2-NuoM harbouring *E. coli* BW29427 into R569, the *E. coli* helper HB101( pRK2013) was used.

Conjugation and selection was performed using gentamycin and nitrofurantoin as selection marker as described before (Ordon et al. 2025).

### Analysis of conservation of NADH dehydrogenase sequences

nuo, ndh, and nqr NADH dehydrogenase amino acid sequences from *Pseudomonas aeruginosa* PAO1 were used for an initial tBLASTn search of the genomes of the culture-collection *Pseudomonas* isolates. for nuo and ndh BLAST hits, the R569 sequences were used in a subsequent tBLASTn. As the BLAST for nqr did not yield any hits for R569, the sequences for IT379 were used for a subsequent tBLASTn. For each gene, the best BLAST hit from each strain was compared to the BLAST query to generate the heat map.

### Bacterial qRT-PCR

Bacterial strains were inoculated in liquid ½ TSB and grown overnight. Cultures were centrifuged to remove the media, and RNA was extracted with TRIzol Reagent. DNA was removed with a DNA-free DNA Removal Kit. SuperScript IV Reverse Transcriptase (Catalog # 18090050, Thermo) was used for the reverse transcription with random oligomer primers. qPCR was performed with Bio-Rad iQ SYBR-green Supermix (Catalog # 1708882). Relative expression for the *nuo* genes was determined with the 2^-ΔΔCt^ method (Livak and Schmittgen 2001), using the geometric mean of the Ct for *recA* and *tpiA* as an internal reference (Wang et al. 2025) and wild-type R569 as the calibrator. Primers used for qRT-PCR are shown in **Supplementary Table 5**.

### QTL mapping

Due to the number of lines, RILs were grown in multiple independent experiments. Within an experiment, the relative root length of each RIL was calculated as the ratio of the mean primary root lengths between the R569 and germ-free treatments. To allow comparison between independent experiments, a Col-0 control was always included and the relative root lengths within each experiment were normalized to a Col-0 relative root length of 1.

The normalized relative root length was used as the phenotype for QTL mapping using the R package R/qtl (Broman et al. 2003). After an initial QTL mapping analysis using previously generated genotype data (Gerald et al. 2014), we generated CAPS markers on chromosome 5 and used these to genotype the RILs (**Supplementary Table 6**). The final QTL mapping included the genotype data from the CAPS markers and excluded genotype data for SNP160_5, SNP161_5, and SNP162_5.

### CRISPR-Cas9 gene editing

Guide RNAs for target genes were designed with CHOPCHOP (Labun et al. 2019), and cloned into pDGE constructs (Ordon et al. 2017). Assembled constructs in pDGE347 (**Supplementary Table 7**) were transformed into *A. thaliana* by floral dipping. Gene edits were identified by PCR amplification and Sanger sequencing (**Supplementary Table 8**).

## Acknowledgements

We thank Shawn Kraemer, Petra Koechner, and Daniëlle de Hoog for technical assistance. We thank Dr. Tak Lee for advice and scripts for data analysis. We thank Dr. Angela Hancock and Dr. Célia Neto for seeds of the *A. thaliana* natural accessions. We thank Cara Haney for the kind gift of the pEXG2 vector.

## Author contributions

CC and PSL conceived of the research, designed experiments, and wrote the manuscript. CC performed most of the experiments and analyzed most of the data. EL performed the bacterial forward genetic screen, generated the R569 Δ*nuoM* mutant, and performed colony count assays. MM and AA generated the IT466 Δ*nuo* mutant. AV and CC together generated the *Pseudomonas* phylogeny and AV assisted with the QTL mapping.

---

**Extended Data Fig. 1.**
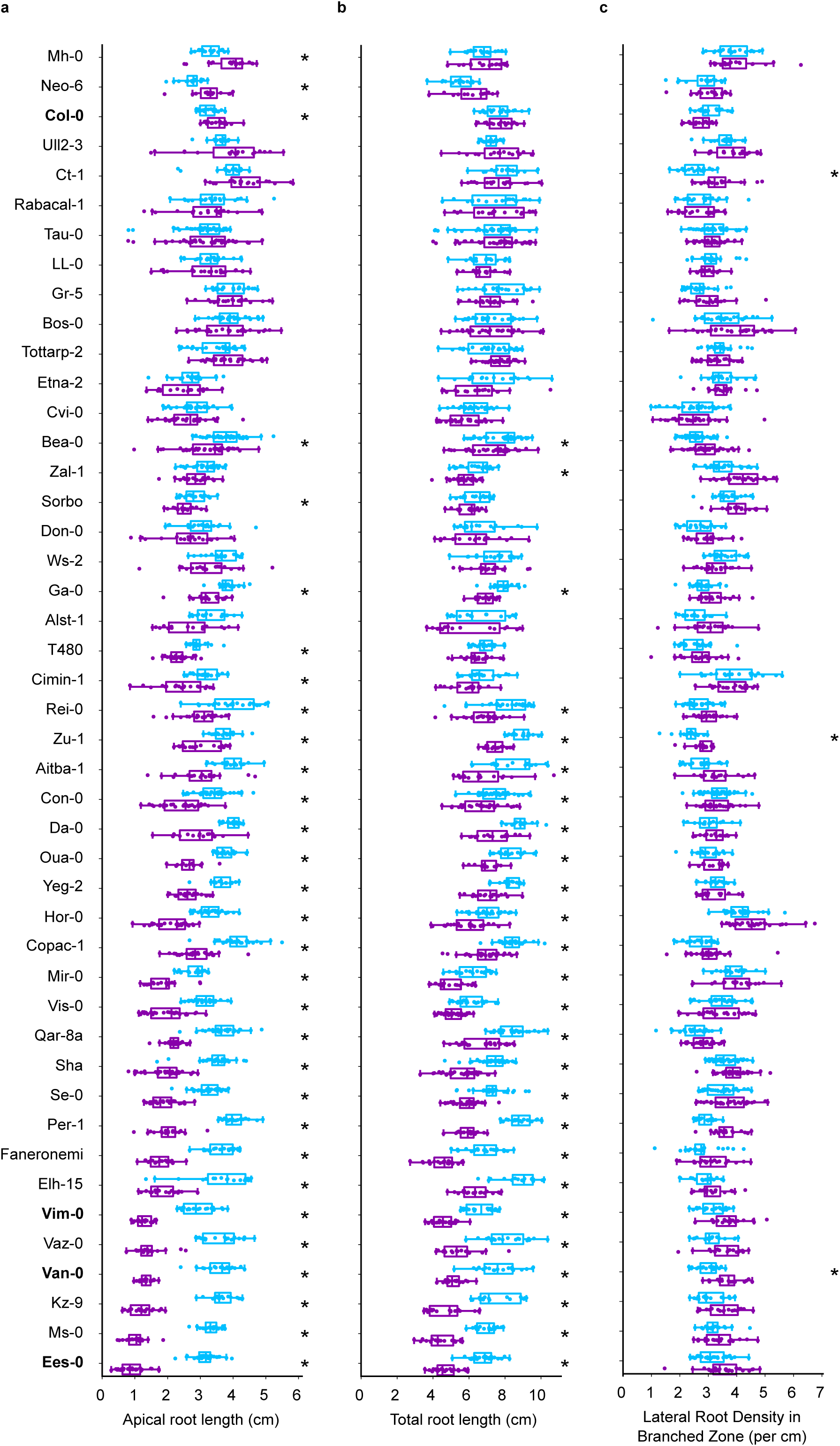
Natural variation in *A. thaliana* root architecture responses to root-associated *Pseudomonas* isolate R569. Apical root length **(a)**, total primary root length **(b)**, and lateral root density in the branched zone **(c)** of 16-day-old *A. thaliana* natural accessions grown in germ-free conditions (light blue) or with *Pseudomonas* R569 (dark purple) for 7 days. In the box plots, boxes represent the interquartile range (IQR) and whiskers represent 1.5 IQR beyond the boxes. The median is shown as a line within the box. Points represent root measurements from individual plants. For each accession, the results of at least 2 independent biological replicates are shown. Asterisks represent a statistically significant difference between the R569-treated and germ-free conditions within each accession, as calculated by pairwise Kruskal-Wallis tests followed by a Benjamini-Hochburg multiple testing correction (false discovery rate = 0.05). Accessions that are further described in this study are highlighted with bold text.

**Extended Data Fig. 2.**
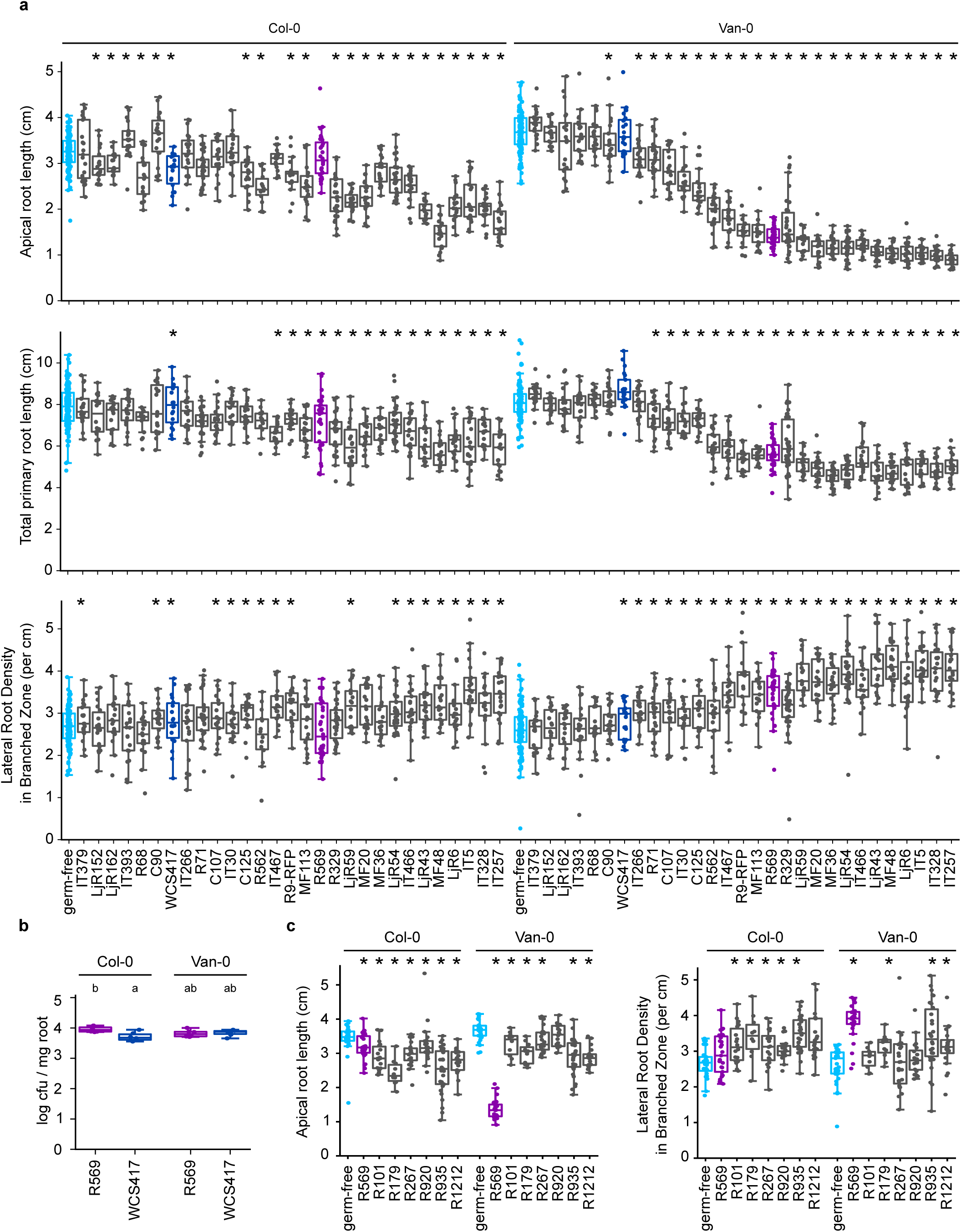
Natural variation in RSA-altering activity among microbiota-derived *Pseudomonas* isolates. **a,** Apical root zone length, total primary root length, and lateral root density in the branched zone of 16-day-old Col-0 and Van-0 grown for 7 days with *Pseudomonas* isolates from microbiota culture collections. Asterisks represent a statistically significant difference for each bacterial treatment compared to the germ-free condition, within each accession, as calculated by pairwise Kruskal-Wallis tests followed by a Benjamini-Hochburg multiple testing correction (false discovery rate = 0.05). For each isolate, the results of at least 2 independent biological replicates are shown. For the germ-free condition, all independent biological replicates are shown; however, only germ-free data from the relevant biological replicates was used for the statistical testing. The germ-free, WCS417, and R569 treatments are highlighted in light blue, dark blue, and dark purple, respectively. **b**, Bacterial load of R569 or WCS417 on roots of 16-day-old Col-0 or Van-0 plants after growth in monoassociation for 7 days. **c,** Apical root zone length and lateral root density in the branched zone of Col-0 and Van-0 grown with bacterial isolates representing core taxa of the root microbiota.

**Extended Data Fig. 3.**
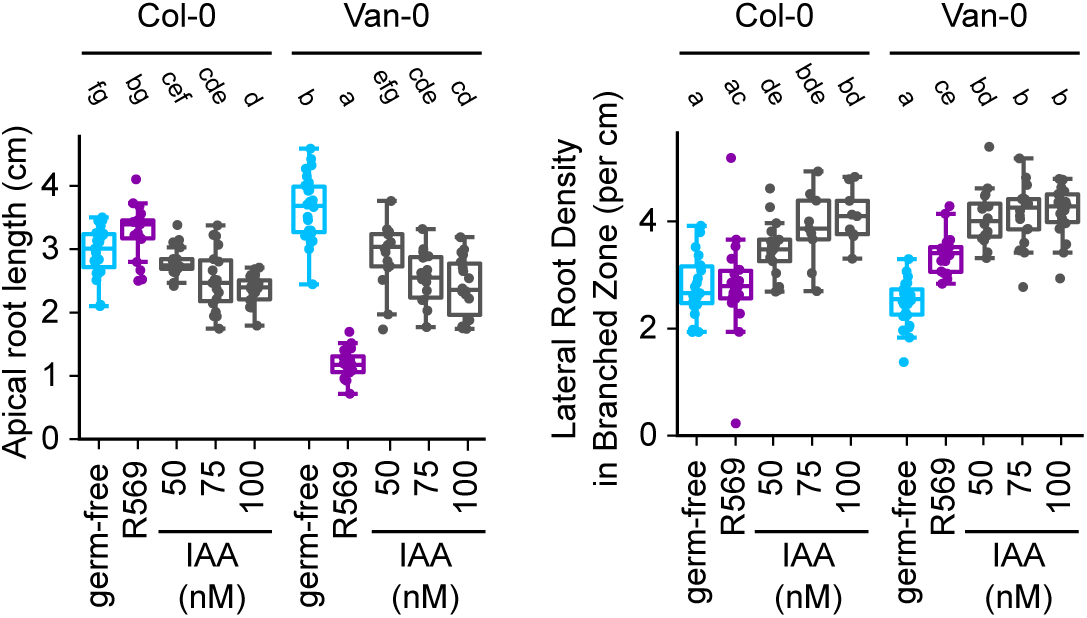
Col-0 and Van-0 show similar root architecture changes in response to exogenous auxin. Apical root length and lateral root density in the branched zone of Col-0 and Van-0 grown with varying concentrations of IAA or in monoassociation with R569.

**Extended Data Fig. 4.**
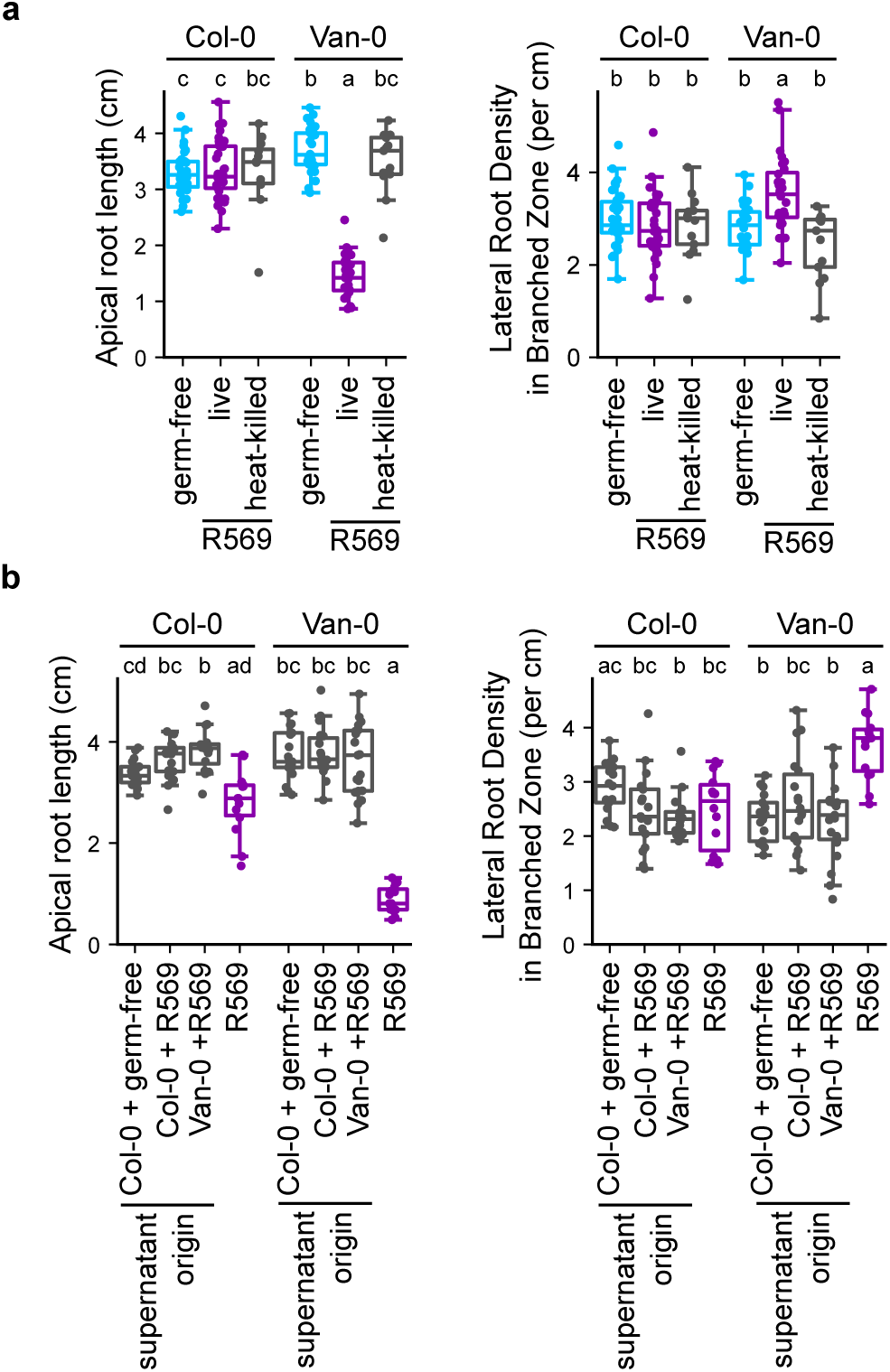
R569-mediated RSA changes require live bacteria in close association with the roots. **a**, Apical root length and lateral root density in the branched zone of Col-0 and Van-0 grown with heat-killed R569. **b**, Apical root length and lateral root density in the branched zone of Col-0 and Van-0 grown on plates prepared with cell-free supernatants originating from R569 cultures grown together with Col-0 or Van-0 seedlings. Cell-free supernatants were prepared by filter sterilizing cultures of the bacterial strains grown in liquid ½ MS medium on plates with the different accessions, indicated by the supernatant origin.

**Extended Data Fig. 5.**
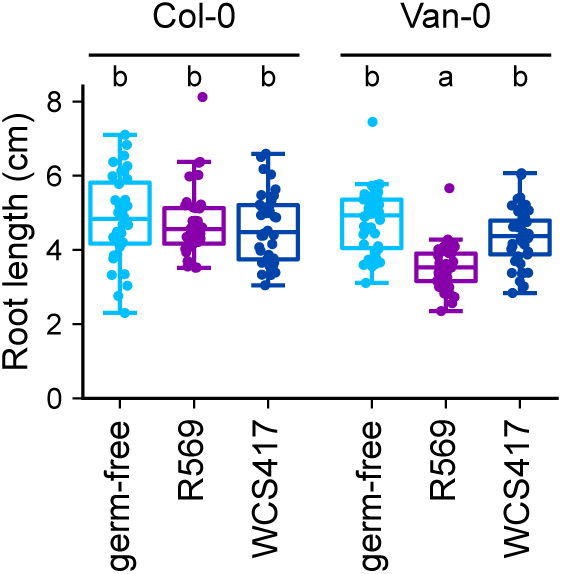
R569-mediated RSA changes are retained in a different gnotobiotic system. Root length of 3-week-old Col-0 and Van-0 grown in perlite with R569, WCS417, or in germ-free conditions.

**Extended Data Fig. 6.**
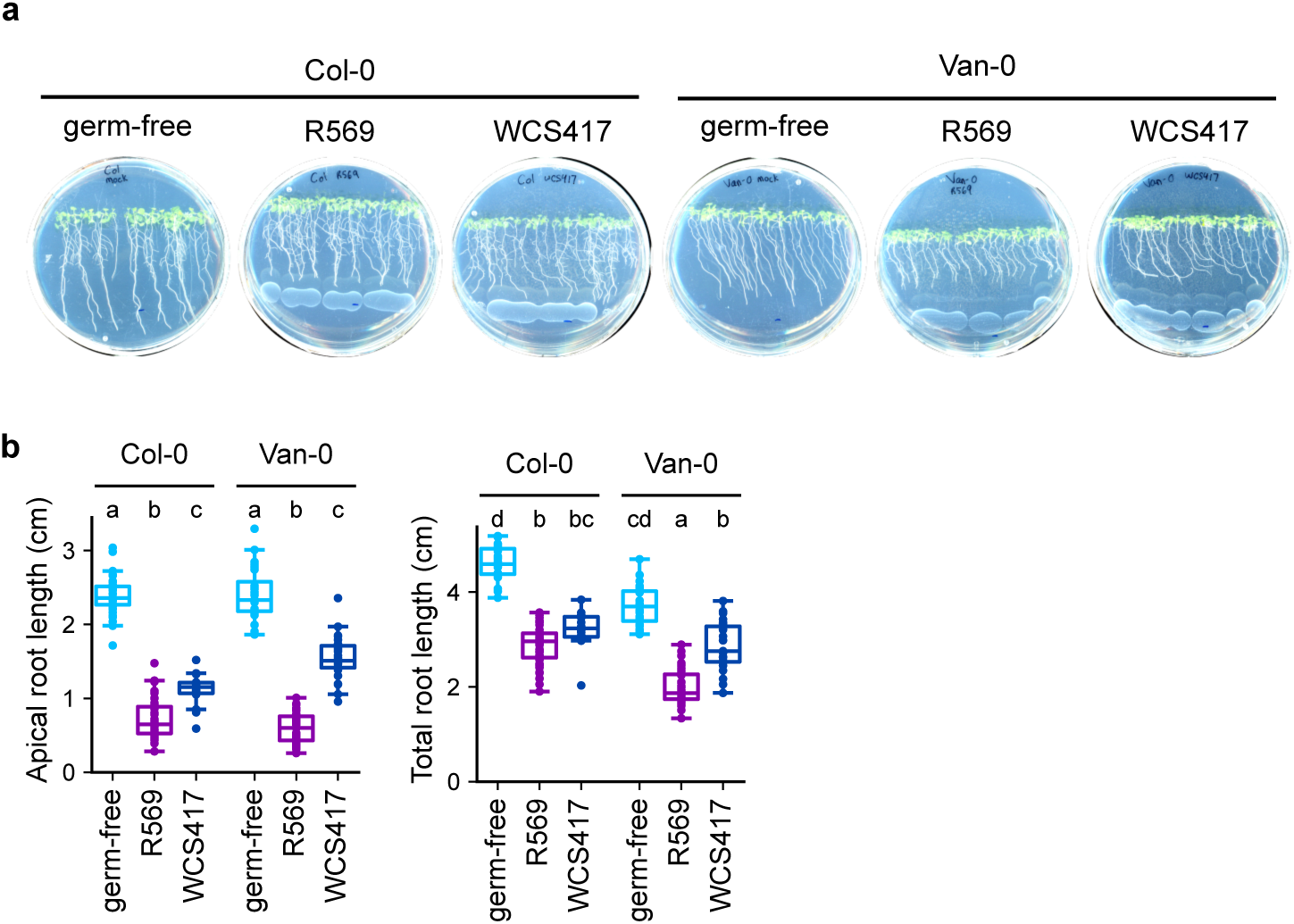
WCS417 causes RSA in both Col-0 and Van-0 when inoculated at a high bacterial density. **a,** Representative images of plates with 12-day-old Col-0 and Van-0 grown with bacteria suspension spotted on the media surface. Bacteria was added 4 days after germination. **b,** Apical root zone length and total primary root length of plants grown as in **(a)**.

**Extended Data Fig. 7.**
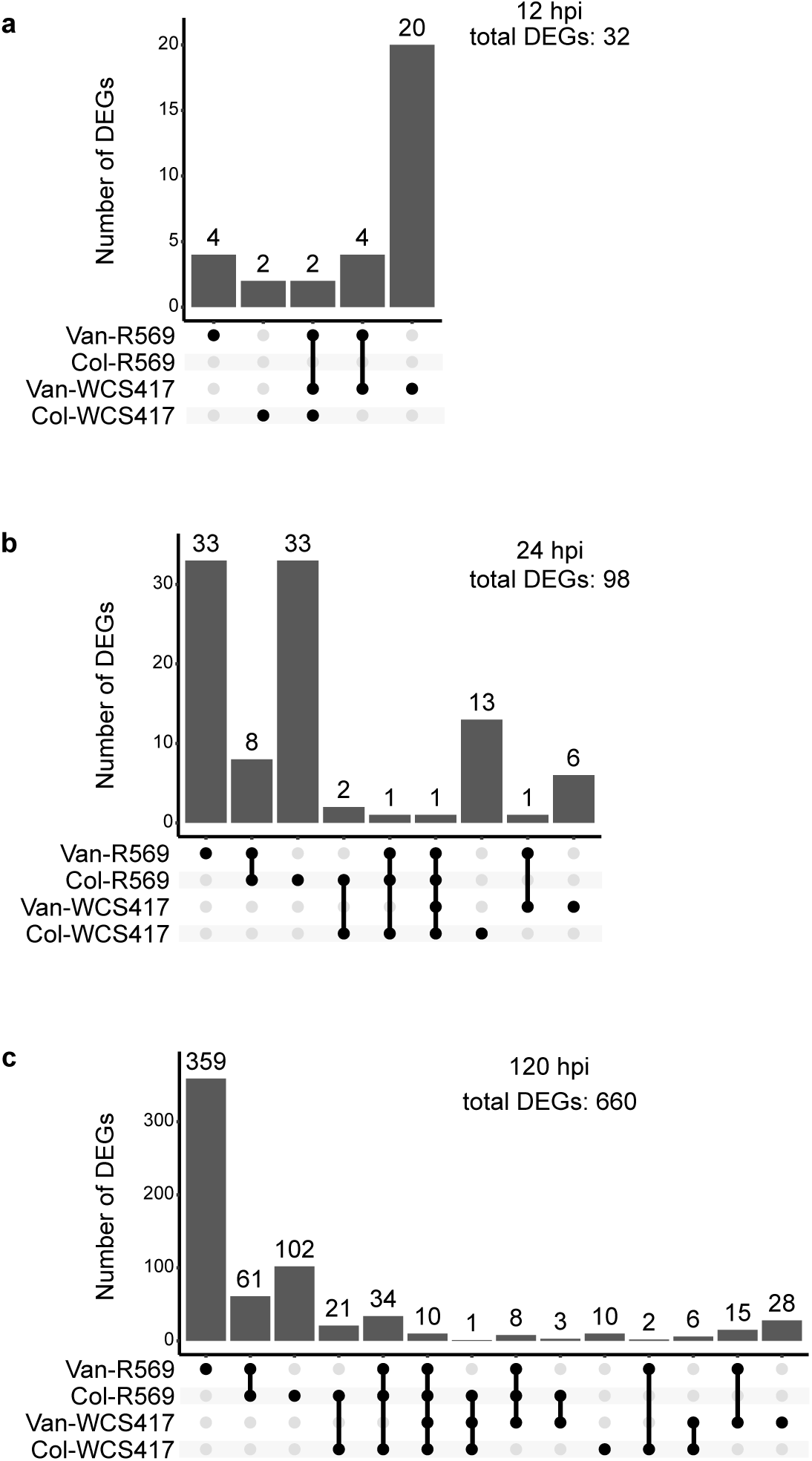
Roots show mild transcriptional responses to bacteria at early time points. **a-c,** Upset plots showing number of root DEGs in plants at 12 h **(a)**, 24 h **(b)**, and 120 h **(c)** hours after inoculation with the indicated bacterial isolates.

**Extended Data Fig. 8.**
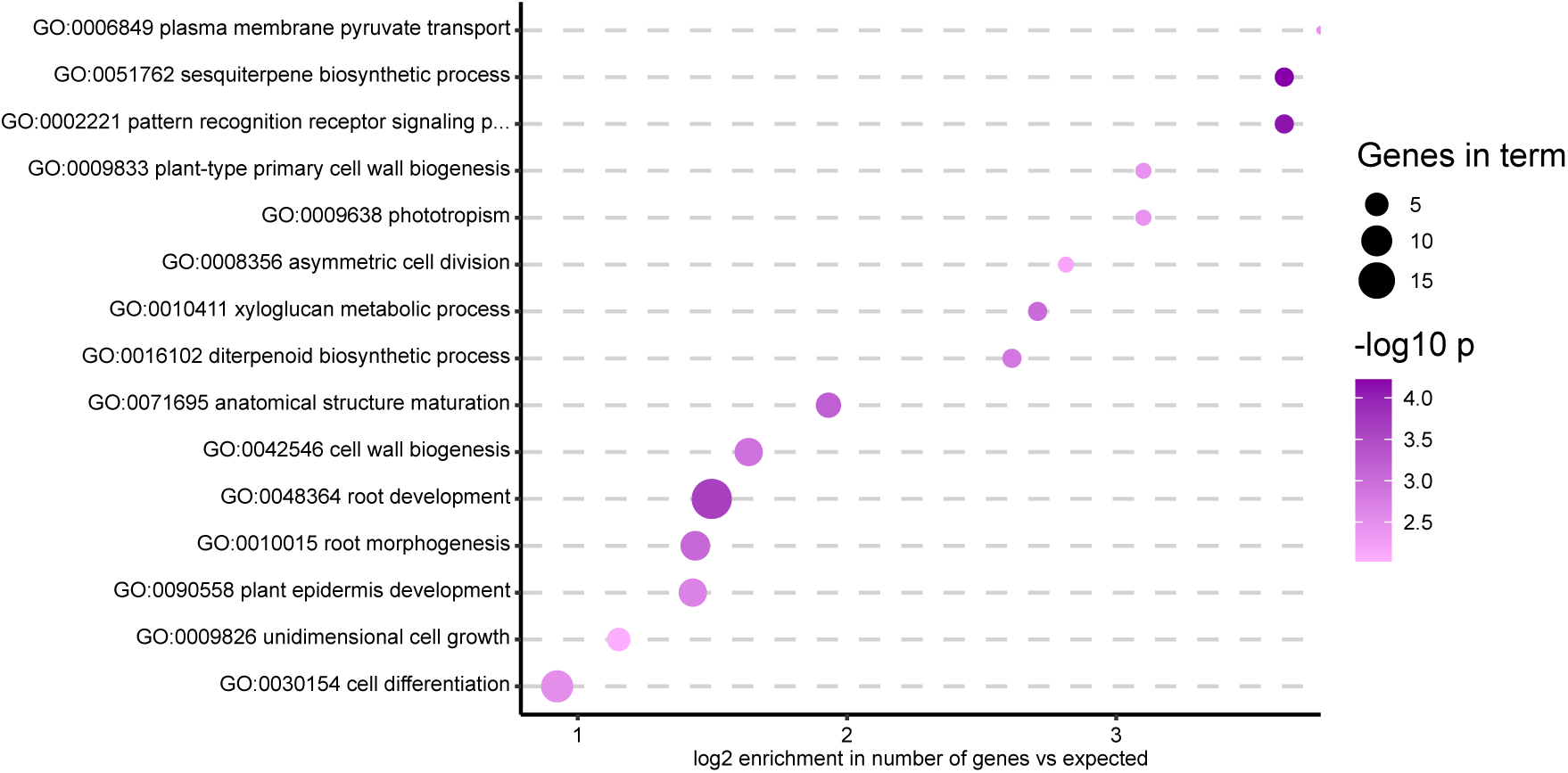
R569 causes accession-specific downregulation of development-related genes in Van-0. GO Biological Process terms enrichment in genes that show significantly stronger transcriptional downregulation by R569 treatment in Van-0 compared to Col-0 at 48 hpi. Significant DEGs were assessed based on the adjusted p < 0.05 for the interaction between genotype and bacterial treatment.

**Extended Data Fig. 9.**
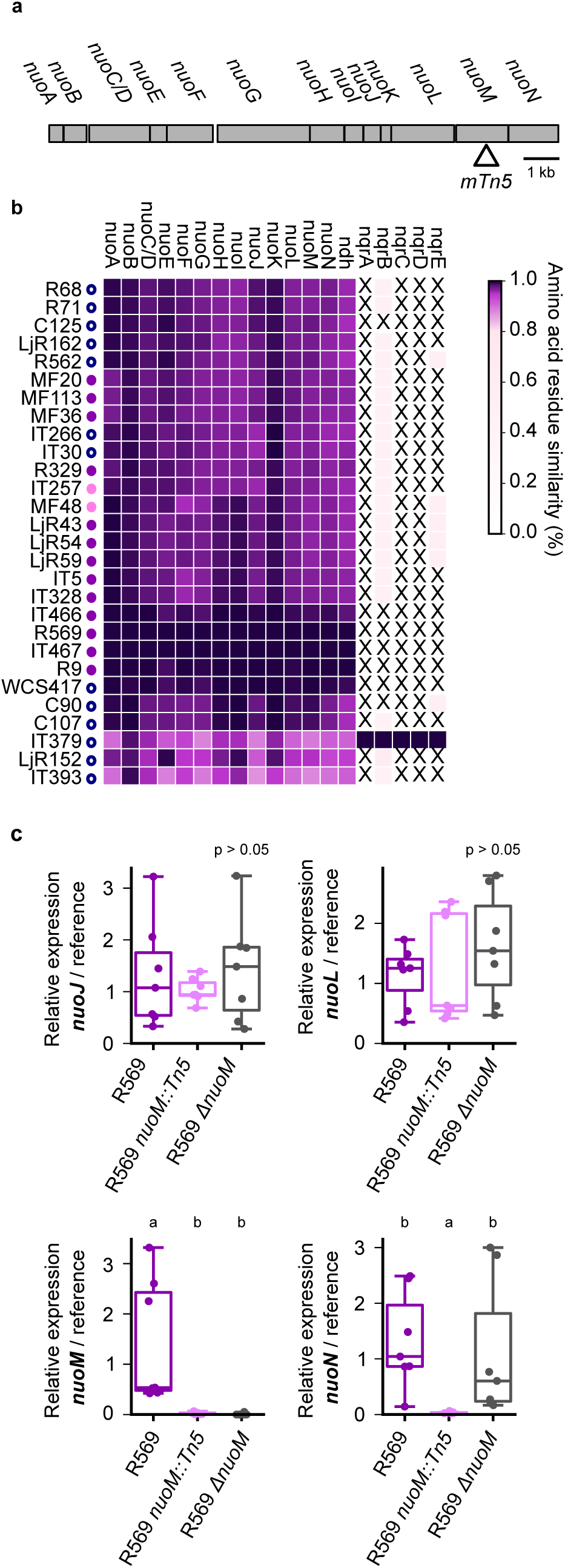
Characterization of the R569 *nuoM* mutants. **a,** Diagram of the *nuo* operon with the location of the *mTn5* insertion in R569 *nuoM::Tn5* marked by an arrowhead. **b,** Heat map showing conservation of NADH dehydrogenases in microbiota-derived *Pseudomonas* isolates, based on percent similarity of the deduced amino acid sequences. Percent similarity was calculated with reference to the R569 for nuo and ndh sequences, and to IT379 for nqr sequences. “X” indicates that no corresponding sequence was found. Circles beside each strain indicate their effect on RSA, as in Fig. 1i. **c,** Expression of *nuo* genes in 12h bacterial cultures of the indicated genotypes, determined with qRT-PCR. *recA* and *tpiA* were used as the reference.

**Extended Data Fig. 10.**
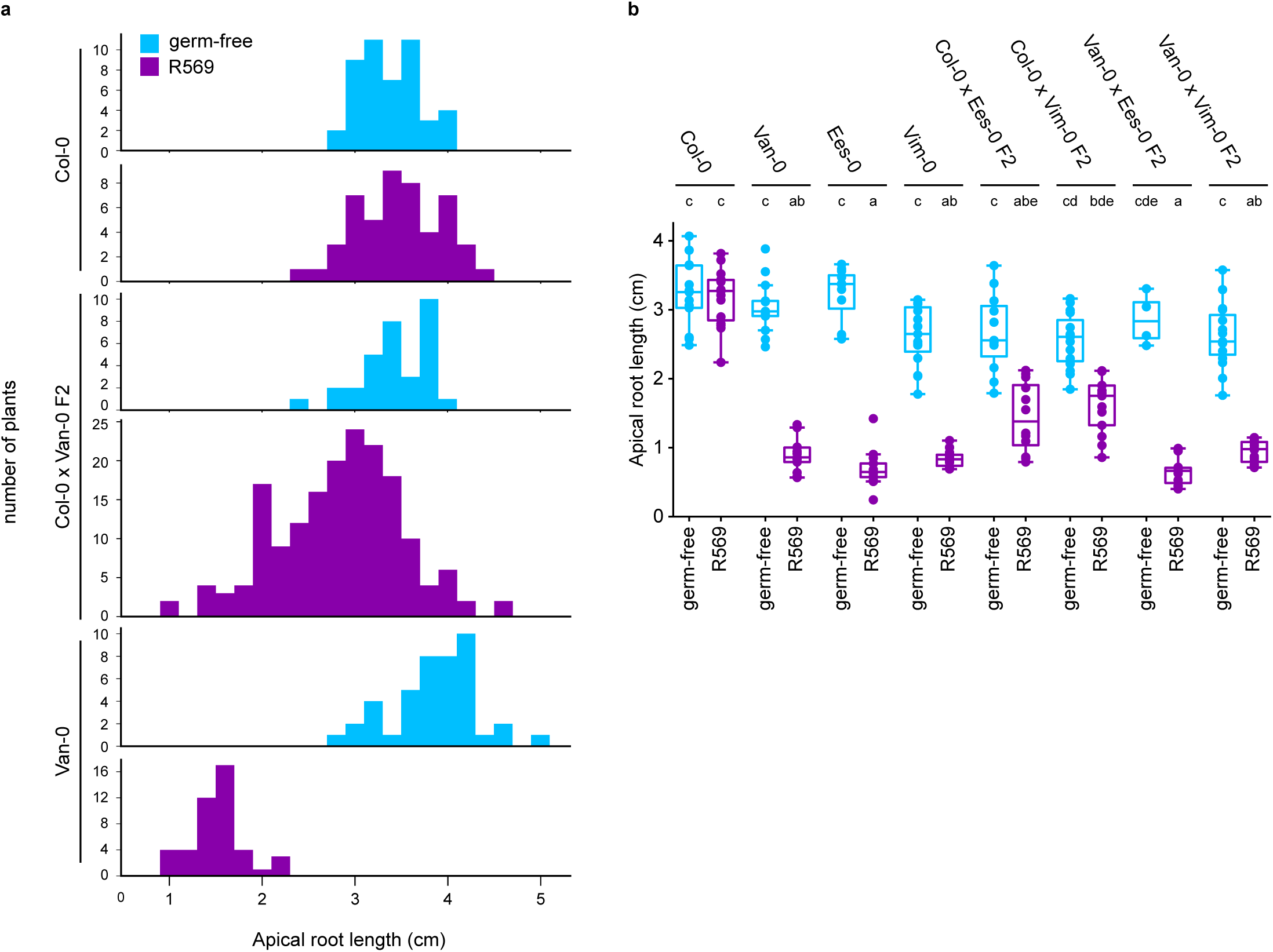
Genetic analysis of accession-specific RSA changes. **a,** Histograms showing the distribution of apical root length of 16-day-old Col-0, Van-0, and F2 offspring of a Col-0 x Van-0 cross, grown in germ-free conditions (light blue) or in monoassociation (dark purple) with R569 for 7 days. **b,** Apical root length of F2 offspring from crosses of Ees-0 and Vim-0 with Col-0 or Van-0, grown in germ-free conditions or in monoassociation with R569.

**Extended Data Fig. 11.**
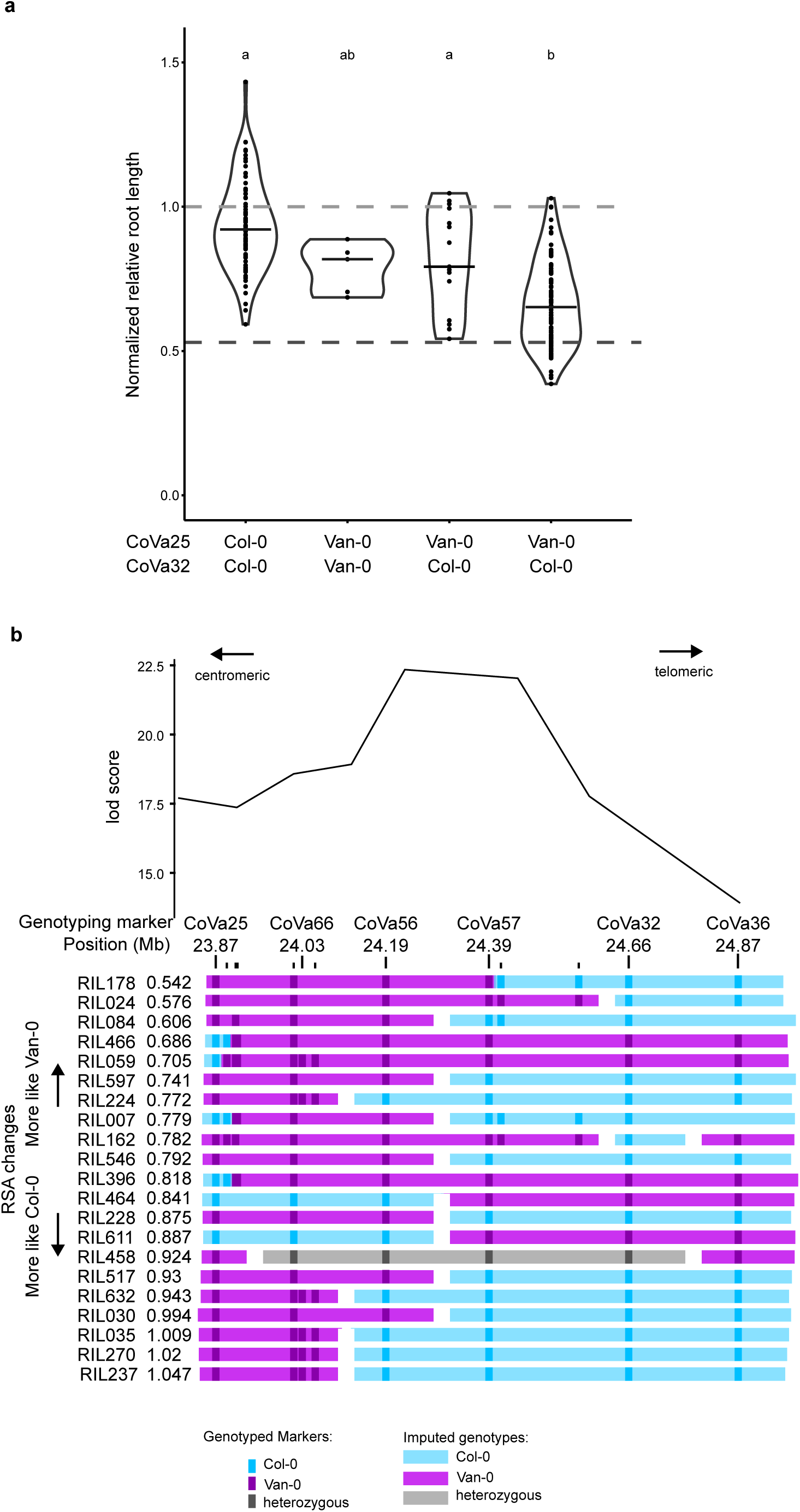
Recombinant inbred lines within the QTL region tend to show a Col-0 root phenotype. **a**, Normalized relative root length of recombinant inbred lines grouped by their genotypes at CoVa25 and CoVa32, two genotyping markers within flanking zones of the QTL region. Each point represents the mean normalized relative root length of multiple individuals from one RIL. The horizontal black line indicates the median for each group. The horizontal dashed lines represent normalized relative root length of Col-0 (light gray) and Van-0 (dark gray). **b,** Graphical genotypes for RILs with recombination break points within the QTL region. The mean normalized relative root length is shown for each RIL.

**Extended Data Fig. 12.**
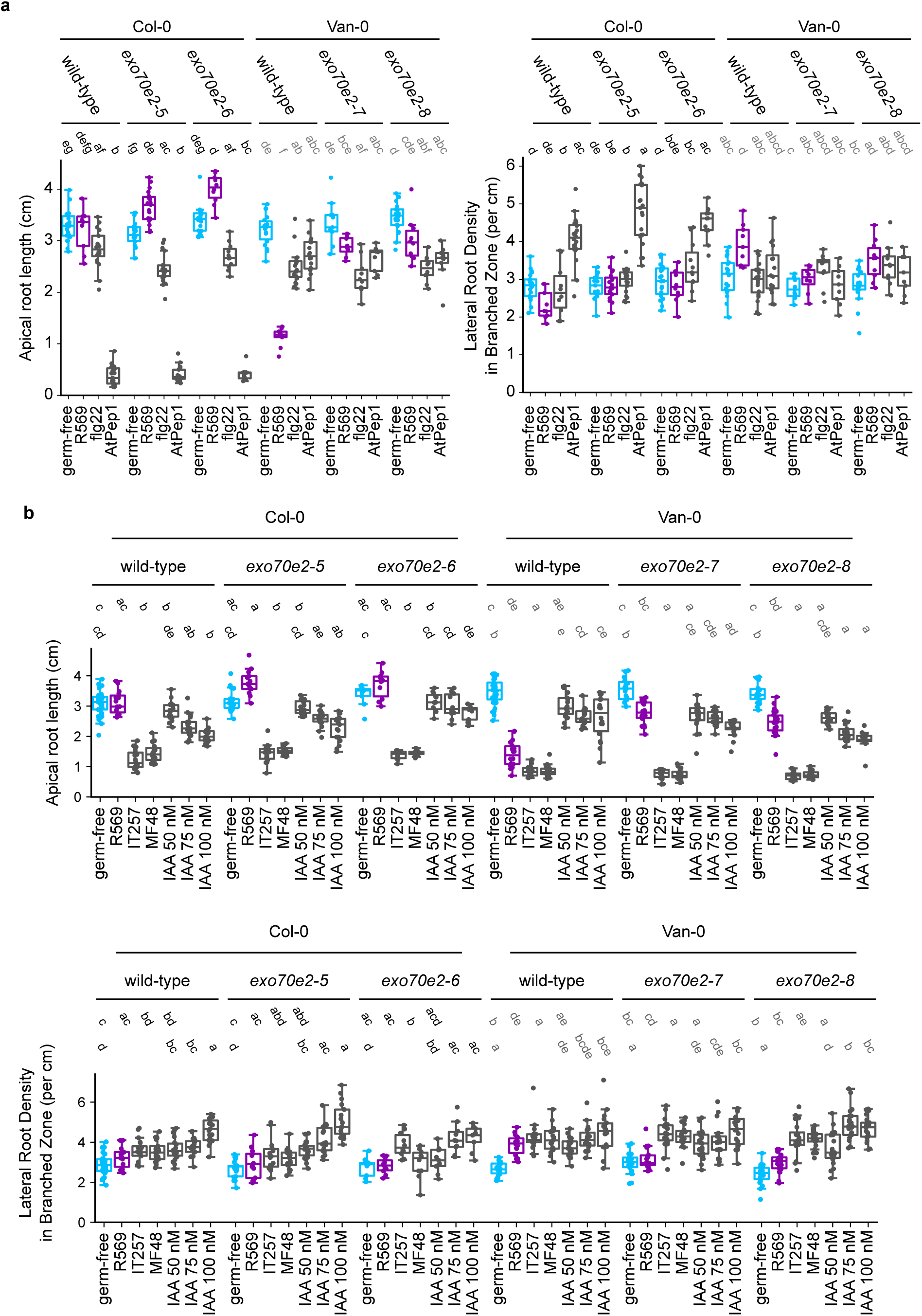
*EXO70E2* is not required for non-accession-specific RSA changes. **a,** Apical root length of *exo70e2* mutants in Col-0 or Van-0 background grown with elicitors flg22 or AtPep1. To avoid an excessive number of statistical comparisons, statistical tests were performed separately for Col-0 and Van-0 backgrounds (statistical letter groups shown in black and grey, respectively). **b,** Apical root length and lateral root density in the branched zone of *exo70e2* mutants in Col-0 or Van-0 background, grown in monoassociation with the indicated *Pseudomonas* isolates or varying concentrations of IAA. Letter groups indicate statistical significance (p < 0.05) according to a Kruskal-Wallis test followed by Dunn’s posthoc test, with a Benjamini-Hochburg posthoc correction. To avoid an excessive number of statistical comparisons, statistical tests were performed separately for bacterial treatments and IAA treatments (statistical letter groups in upper and lower rows, respectively), and for Col-0 and Van-0 backgrounds (statistical letter groups shown in black and grey, respectively).

**Extended Data Fig. 13.**
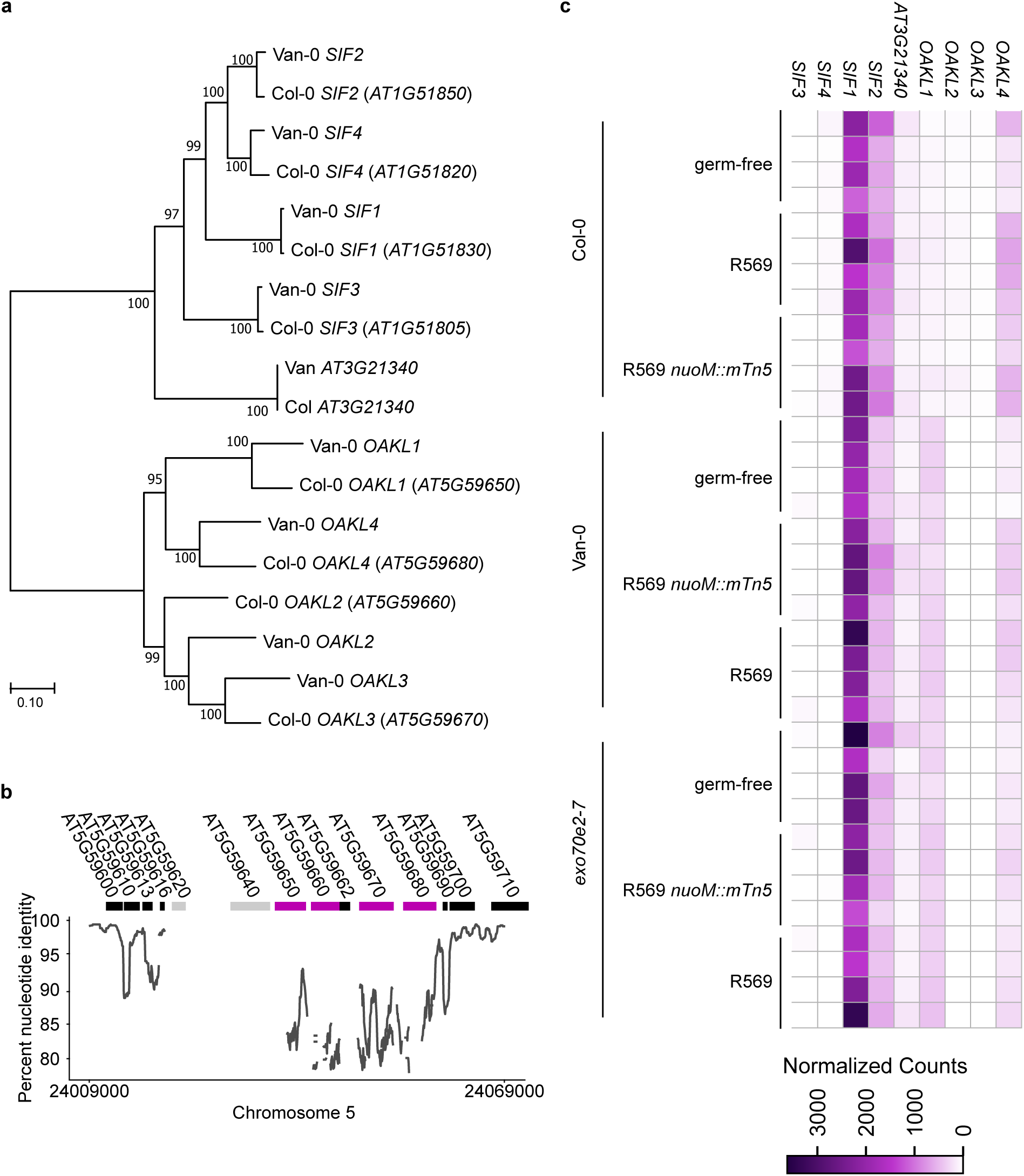
Divergence of the *OAKL* cluster in Col-0 and Van-0. **a,** Maximum-likelihood phylogeny of *OAKL* genes and closely related MLD-LRR-RLKs. Van-0 *OAKL* and *SIF* genes were named according to their physical position within the cluster. Numbers at nodes represent bootstrap consensus support based on 500 replicates. **b,** Similarity between Col-0 and Van-0 genomic sequence in the *OAKL* cluster and flanking regions. Percent nucleotide identity was calculated in 500 bp sliding windows. Discontinuities in the line represent regions that could not be aligned. Genetic loci are shown above the plot: *OAKL* genes are highlighted in purple, pseudogenes or transposable elements are shown in grey, and other genes are in black. Genomic coordinates are shown according to TAIR10. Note that the scale on the y-axis does not begin at 0. **c,** Heat map showing expression of MLD-LRR-RLK genes of the *OAKL* and *SIF* families. Counts are from the RNA-seq experiment presented in Fig. 5 and normalized to account for differences in coverage between samples.

**Extended Data Fig. 14.**
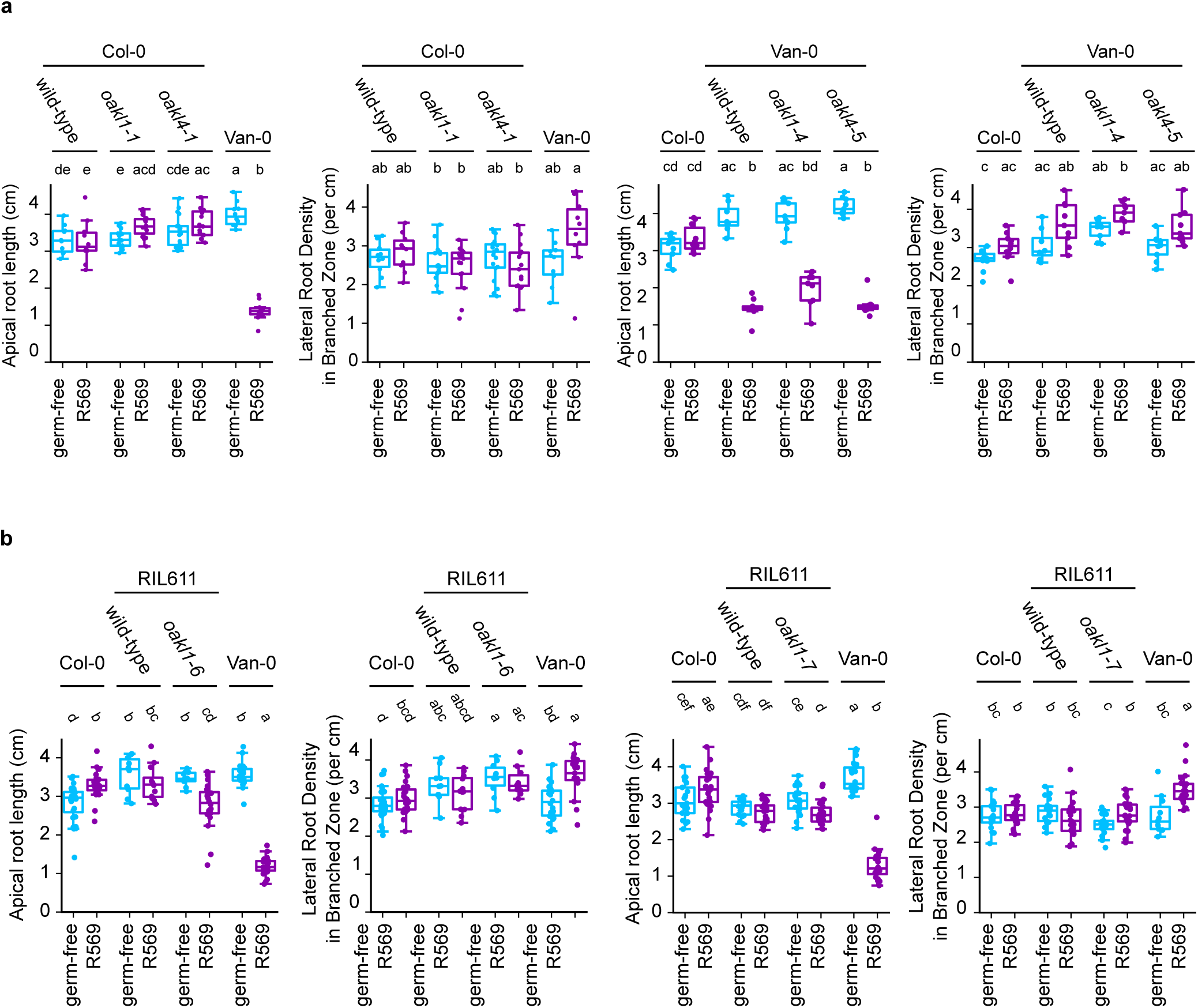
Gene editing of *OAKL1* individually is not sufficient to alter the accession-specific *Pseudomonas*-mediated RSA changes. **a,** Apical root length and lateral root density in the branched zone of *oakl* single mutants in the Col-0 or Van-0 background. **b,** Apical root length and lateral root density in the branched zone of *oakl1* single mutants in the RIL611 background.

**Supplementary Table 1. Frequency of *OAKL1* and *OAKL4* gene edits.** Number of T1 plants with gene edits in the indicated genes. *Due to the sufficient number of gene-edited lines recovered, not all T1s were analyzed by sequencing; some T1s classified with as wild-type based on PCR amplicon size may therefore contain undetected small insertion or deletion gene edits.

**Supplementary Table 2. SynCom composition**

**Supplementary Table 3. Culture collections used in this study.**

**Supplementary Table 4. Primers used for cloning**

**Supplementary Table 5. Primers used for qRT-PCR**

**Supplementary Table 6. CAPS markers used for genotyping RILs.** Positions of the polymorphisms are given according to TAIR10. The reference and alternate band sizes refer to the size of the expected DNA fragment after PCR amplification and restriction digestions of the Col-0 and Van-0 alleles, respectively.

**Supplementary Table 7. sgRNAs used for crispr-Cas9 gene editing**

**Supplementary Table 8. Primers used for genotyping gene-edited lines**

**Supplementary Dataset 1. DESeq2 output for RNA-seq experiments. a,b,** DESeq2 output for the 48 hpi time point from the RNA-seq experiment shown in Fig. 2, showing log2 fold changes (log2FC) and adjusted p values (padj) for the given comparisons, showing all analyzed genes **(a)** or genes with significant expression differences in at least one comparison **(b)**. **c,d,** DESeq2 output for the RNA-seq experiment shown in Fig. 5, showing log2 fold changes (log2FC) and adjusted p values (padj) for the given comparisons, showing all analyzed genes **(a)** or genes with significant expression differences in at least one comparison **(b)**. **e,** DEseq2 output for DESeq2 output for the RNA-seq experiment shown in Fig. 5, using the interaction formula ∼ plant_mutation + bacteria_mutation + plant_mutation:bacteria_mutation to identify significant interactions between the effect of *exo70e2-7* (plant_mutation) and R569 *nuoM::Tn5* (bacteria_mutation).

## References

Alexa A and Rahnenfuhrer J. topGO: Enrichment Analysis for Gene Ontology. 2025. doi:10.18129/B9.bioc.topGO

Bai Y, Müller DB, Srinivas G, Garrido-Oter R, Potthoff E, Rott M, Dombrowski N, Münch PC, Spaepen S, Remus-Emsermann M, et al. Functional overlap of the Arabidopsis leaf and root microbiota. Nature. 2015:528(7582):364–369. 10.1038/nature16192

Banerjee S and van der Heijden MGA. Soil microbiomes and one health. Nat Rev Microbiol. 2023:21(1):6–20. 10.1038/s41579-022-00779-w

Bouain N, Korte A, Satbhai SB, Nam H-I, Rhee SY, Busch W, and Rouached H. Systems genomics approaches provide new insights into Arabidopsis thaliana root growth regulation under combinatorial mineral nutrient limitation. PLOS Genetics. 2019:15(11):e1008392. 10.1371/journal.pgen.1008392

Broman KW, Wu H, Sen S, and Churchill GA. R/qtl: QTL mapping in experimental crosses. Bioinformatics. 2003:19(7):889–890. 10.1093/bioinformatics/btg112

Capilla-Pérez L, Solier V, Gilbault E, Lian Q, Goel M, Huettel B, Keurentjes JJB, Loudet O, and Mercier R. Enhanced recombination empowers the detection and mapping of Quantitative Trait Loci. Commun Biol. 2024:7(1):829. 10.1038/s42003-024-06530-w

Chan C, Panzeri D, Okuma E, Tõldsepp K, Wang Y-Y, Louh G-Y, Chin T-C, Yeh Y-H, Yeh H-L, Yekondi S, et al. STRESS INDUCED FACTOR 2 Regulates Arabidopsis Stomatal Immunity through Phosphorylation of the Anion Channel SLAC1. Plant Cell. 2020:32(7):2216–2236. 10.1105/tpc.19.00578

Chen S. Ultrafast one-pass FASTQ data preprocessing, quality control, and deduplication using fastp. iMeta. 2023:2(2):e107. 10.1002/imt2.107

Ciemniecki JA and Newman DK. NADH dehydrogenases are the predominant phenazine reductases in the electron transport chain of Pseudomonas aeruginosa. Molecular Microbiology. 2023:119(5):560–573. 10.1111/mmi.15049

Conway JM, Walton WG, Salas-González I, Law TF, Lindberg CA, Crook LE, Kosina SM, Fitzpatrick CR, Lietzan AD, Northen TR, et al. Diverse MarR bacterial regulators of auxin catabolism in the plant microbiome. Nat Microbiol. 2022:7(11):1817–1833. 10.1038/s41564-022-01244-3

Copeland C, Schulze-Lefert P, and Ma K-W. Potential and challenges for application of microbiomes in agriculture. The Plant Cell. 2025:koaf185. 10.1093/plcell/koaf185

De la Concepcion JC. The exocyst complex is an evolutionary battleground in plant-microbe interactions. Current Opinion in Plant Biology. 2023:76:102482. 10.1016/j.pbi.2023.102482

Deja-Muylle A, Opdenacker D, Parizot B, Motte H, Lobet G, Storme V, Clauw P, Njo M, and Beeckman T. Genetic Variability of Arabidopsis thaliana Mature Root System Architecture and Genome-Wide Association Study. Frontiers in Plant Science. 2022:12(January):1–16. 10.3389/fpls.2021.814110

Ding Y, Wang J, Chun Lai JH, Ling Chan VH, Wang X, Cai Y, Tan X, Bao Y, Xia J, Robinson DG, et al. Exo70E2 is essential for exocyst subunit recruitment and EXPO formation in both plants and animals. Mol Biol Cell. 2014:25(3):412–426. 10.1091/mbc.E13-10-0586

Dini-Andreote F, Wells DM, Atkinson JA, Atkinson BS, Finkel OM, and Castrillo G. Microbial drivers of root plasticity. New Phytologist. 2025:n/a(n/a). 10.1111/nph.70371

Durán P, Flores-Uribe J, Wippel K, Zhang P, Guan R, Melkonian B, Melkonian M, and Garrido-Oter R. Shared features and reciprocal complementation of the Chlamydomonas and Arabidopsis microbiota. Nat Commun. 2022:13(1):406. 10.1038/s41467-022-28055-8

Fernández-Calvo P, López G, Martín-Dacal M, Aitouguinane M, Carrasco-López C, González-Bodí S, Bacete L, Mélida H, Sánchez-Vallet A, and Molina A. Leucine rich repeat-malectin receptor kinases IGP1/CORK1, IGP3 and IGP4 are required for arabidopsis immune responses triggered by β-1,4-D-Xylo-oligosaccharides from plant cell walls. The Cell Surface. 2024:11:100124. 10.1016/j.tcsw.2024.100124

Finkel OM, Salas-González I, Castrillo G, Conway JM, Law TF, Teixeira PJPL, Wilson ED, Fitzpatrick CR, Jones CD, and Dangl JL. A single bacterial genus maintains root growth in a complex microbiome. Nature. 2020:587(7832):103–108. 10.1038/s41586-020-2778-7

Gerald JNF, Carlson AL, Smith E, Maloof JN, Weigel D, Chory J, Borevitz JO, and Swanson RJ. New Arabidopsis advanced intercross recombinant inbred lines reveal female control of nonrandom mating. Plant Physiology. 2014:165(1):175–185. 10.1104/pp.113.233213

Getzke F, Hassani MA, Crüsemann M, Malisic M, Zhang P, Ishigaki Y, Böhringer N, Jiménez Fernández A, Wang L, Ordon J, et al. Cofunctioning of bacterial exometabolites drives root microbiota establishment. Proceedings of the National Academy of Sciences. 2023:120(15):e2221508120. 10.1073/pnas.2221508120

Gonin M, Salas-González I, Gopaulchan D, Frene JP, Roden S, Van de Poel B, Salt DE, and Castrillo G. Plant microbiota controls an alternative root branching regulatory mechanism in plants. Proc Natl Acad Sci USA. 2023:120(15):e2301054120. 10.1073/pnas.2301054120

Holden S, Bergum M, Green P, Bettgenhaeuser J, Hernández-Pinzón I, Thind A, Clare S, Russell JM, Hubbard A, Taylor J, et al. A lineage-specific Exo70 is required for receptor kinase–mediated immunity in barley. Science Advances. 2022:8(27):eabn7258. 10.1126/sciadv.abn7258

Hreha TN, Foreman S, Duran-Pinedo A, Morris AR, Diaz-Rodriguez P, Jones JA, Ferrara K, Bourges A, Rodriguez L, Koffas MAG, et al. The three NADH dehydrogenases of Pseudomonas aeruginosa: Their roles in energy metabolism and links to virulence. PLOS ONE. 2021:16(2):e0244142. 10.1371/journal.pone.0244142

Huebbers JW, Caldarescu GA, Kubátová Z, Sabol P, Levecque SCJ, Kuhn H, Kulich I, Reinstädler A, Büttgen K, Manga-Robles A, et al. Interplay of EXO70 and MLO proteins modulates trichome cell wall composition and susceptibility to powdery mildew. Plant Cell. 2024:36(4):1007– 1035. 10.1093/plcell/koad319

Kellermeier F, Armengaud P, Seditas TJ, Danku J, Salt DE, and Amtmann A. Analysis of the Root System Architecture of *Arabidopsis* Provides a Quantitative Readout of Crosstalk between Nutritional Signals. The Plant Cell. 2014:26(4):1480–1496. 10.1105/tpc.113.122101

Keppler A, Roulier M, Pfeilmeier S, Petti GC, Sintsova A, Maier BA, Bortfeld-Miller M, Sunagawa S, Zipfel C, and Vorholt JA. Plant microbiota feedbacks through dose-responsive expression of general non-self response genes. Nat Plants. 2025:11(1):74–89. 10.1038/s41477-024-01856-z

Kim D, Paggi JM, Park C, Bennett C, and Salzberg SL. Graph-based genome alignment and genotyping with HISAT2 and HISAT-genotype. Nat Biotechnol. 2019:37(8):907–915. 10.1038/s41587-019-0201-4

Koch BL, Gardner D, Smith H, Bracewell R, Awdey L, Foster J, Borniego ML, Munch DH, Nielsen ME, Pasupuleti R, et al. Molecular Insights into the Production of Extracellular Vesicles by Plants. 2025:2025.06.16.659989. 10.1101/2025.06.16.659989

Kolmogorov M, Yuan J, Lin Y, and Pevzner PA. Assembly of long, error-prone reads using repeat graphs. Nat Biotechnol. 2019:37(5):540–546. 10.1038/s41587-019-0072-8

Labun K, Montague TG, Krause M, Torres Cleuren YN, Tjeldnes H, and Valen E. CHOPCHOP v3: expanding the CRISPR web toolbox beyond genome editing. Nucleic Acids Res. 2019:47(W1):W171–W174. 10.1093/nar/gkz365

Lachowiec J, Shen X, Queitsch C, and Carlborg Ö. A Genome-Wide Association Analysis Reveals Epistatic Cancellation of Additive Genetic Variance for Root Length in Arabidopsis thaliana. PLoS Genetics. 2015:11(9):1–21. 10.1371/journal.pgen.1005541

Lamers J, Schippers B, and Geels F. Soil-borne diseases of wheat in the Netherlands and results of seed bacterization with pseudomonads against Gaeumannomyces graminis var. tritici, associated with disease resistance. Cereal breeding related to integrated cereal production. 1988:134–139.

LaRue T, Lindner H, Srinivas A, Exposito-Alonso M, Lobet G, and Dinneny JR. Uncovering natural variation in root system architecture and growth dynamics using a robotics-assisted phenomics platform. eLife. 2022:11:e76968. 10.7554/eLife.76968

Levy A, Salas Gonzalez I, Mittelviefhaus M, Clingenpeel S, Herrera Paredes S, Miao J, Wang K, Devescovi G, Stillman K, Monteiro F, et al. Genomic features of bacterial adaptation to plants. Nature Genetics. 2018:50(1):138–150. 10.1038/s41588-017-0012-9

Liao Y, Smyth GK, and Shi W. featureCounts: an efficient general purpose program for assigning sequence reads to genomic features. Bioinformatics. 2014:30(7):923–930. 10.1093/bioinformatics/btt656

Liu J, Wang T, Qin Q, Yu X, Yang S, Dinkins RD, Kuczmog A, Putnoky P, Muszyński A, Griffitts JS, et al. Paired *Medicago* receptors mediate broad-spectrum resistance to nodulation by *Sinorhizobium meliloti* carrying a species-specific gene. Proc Natl Acad Sci USA. 2022:119(51):e2214703119. 10.1073/pnas.2214703119

Liu M-CJ, Yeh F-LJ, Yvon R, Simpson K, Jordan S, Chambers J, Wu H-M, and Cheung AY. Extracellular pectin-RALF phase separation mediates FERONIA global signaling function. Cell. 2024:187(2):312–330.e22. 10.1016/j.cell.2023.11.038

Livak KJ and Schmittgen TD. Analysis of Relative Gene Expression Data Using Real-Time Quantitative PCR and the 2−ΔΔCT Method. Methods. 2001:25(4):402–408. 10.1006/meth.2001.1262

Love MI, Huber W, and Anders S. Moderated estimation of fold change and dispersion for RNA-seq data with DESeq2. Genome Biology. 2014:15(12):550. 10.1186/s13059-014-0550-8

Ma KW, Niu Y, Jia Y, Ordon J, Copeland C, Emonet A, Geldner N, Guan R, Stolze SC, Nakagami H, et al. Coordination of microbe–host homeostasis by crosstalk with plant innate immunity. Nature Plants. 2021:7(6):814–825. 10.1038/s41477-021-00920-2

Ma K-W, Ordon J, and Schulze-Lefert P. Gnotobiotic Plant Systems for Reconstitution and Functional Studies of the Root Microbiota. Current Protocols. 2022:2(2):e362. 10.1002/cpz1.362

Maier BA, Kiefer P, Field CM, Hemmerle L, Bortfeld-miller M, Emmenegger B, Schäfer M, Pfeilmeier S, Sunagawa S, Vogel CM, et al. A general non-self response as part of plant immunity. Nature Plants. 2021:7(May). 10.1038/s41477-021-00913-1

Malivert A and Hamant O. Why is FERONIA pleiotropic? Nat Plants. 2023:9(7):1018–1025. 10.1038/s41477-023-01434-9

Marçais G, Delcher AL, Phillippy AM, Coston R, Salzberg SL, and Zimin A. MUMmer4: A fast and versatile genome alignment system. PLOS Computational Biology. 2018:14(1):e1005944. 10.1371/journal.pcbi.1005944

Martín-Dacal M, Fernández-Calvo P, Jiménez-Sandoval P, López G, Garrido-Arandía M, Rebaque D, del Hierro I, Berlanga DJ, Torres MÁ, Kumar V, et al. Arabidopsis immune responses triggered by cellulose- and mixed-linked glucan-derived oligosaccharides require a group of leucine-rich repeat malectin receptor kinases. The Plant Journal. 2023:113(4):833–850. 10.1111/tpj.16088

Meier M, Liu Y, Lay-Pruitt KS, Takahashi H, and von Wirén N. Auxin-mediated root branching is determined by the form of available nitrogen. Nature Plants. 2020:6(9):1136–1145. 10.1038/s41477-020-00756-2

Melnyk RA, Hossain SS, and Haney CH. Convergent gain and loss of genomic islands drive lifestyle changes in plant-associated Pseudomonas. ISME J. 2019:13(6):1575–1588. 10.1038/s41396-019-0372-5

Northen TR, Kleiner M, Torres M, Kovács ÁT, Nicolaisen MH, Krzyżanowska DM, Sharma S, Lund G, Jelsbak L, Baars O, et al. Community standards and future opportunities for synthetic communities in plant–microbiota research. Nat Microbiol. 2024:9(11):2774–2784. 10.1038/s41564-024-01833-4

Ordon J, Gantner J, Kemna J, Schwalgun L, Reschke M, Streubel J, Boch J, and Stuttmann J. Generation of chromosomal deletions in dicotyledonous plants employing a user-friendly genome editing toolkit. The Plant Journal. 2017:89(1):155–168. 10.1111/tpj.13319

Ordon J, Logemann E, Maier L-P, Lee T, Dahms E, Oosterwijk A, Flores-Uribe J, Miyauchi S, Paoli L, Stolze SC, et al. Conserved immunomodulation and variation in host association by Xanthomonadales commensals in Arabidopsis root microbiota. Nat Plants. 2025:1–20. 10.1038/s41477-025-01918-w

Ortiz-Morea FA, Liu J, Shan L, and He P. Malectin-like receptor kinases as protector deities in plant immunity. Nat Plants. 2021:8(1):27–37. 10.1038/s41477-021-01028-3

Poncini L, Wyrsch I, Tendon VD, Vorley T, Boller T, Geldner N, Métraux JP, and Lehmann S. In roots of Arabidopsis thaliana, the damage-associated molecular pattern AtPep1 is a stronger elicitor of immune signalling than flg22 or the chitin heptamer. PLoS ONE. 2017:12(10):1–21. 10.1371/journal.pone.0185808

Poulsen CP, Dilokpimol A, Mouille G, Burow M, and Geshi N. Arabinogalactan Glycosyltransferases Target to a Unique Subcellular Compartment That May Function in Unconventional Secretion in Plants. Traffic. 2014:15(11):1219–1234. 10.1111/tra.12203

Price MN, Dehal PS, and Arkin AP. FastTree: Computing Large Minimum Evolution Trees with Profiles instead of a Distance Matrix. Mol Biol Evol. 2009:26(7):1641–1650. 10.1093/molbev/msp077

Redditt TJ, Chung E-H, Karimi HZ, Rodibaugh N, Zhang Y, Trinidad JC, Kim JH, Zhou Q, Shen M, Dangl JL, et al. AvrRpm1 Functions as an ADP-Ribosyl Transferase to Modify NOI Domain-Containing Proteins, Including Arabidopsis and Soybean RPM1-Interacting Protein4. The Plant Cell. 2019:31(11):2664–2681. 10.1105/tpc.19.00020

Roux F, Frachon L, and Bartoli C. The Genetic Architecture of Adaptation to Leaf and Root Bacterial Microbiota in Arabidopsis thaliana. Mol Biol Evol. 2023:40(5):msad093. 10.1093/molbev/msad093

Scharwies JD, Clarke T, Zheng Z, Dinneny A, Birkeland S, Veltman MA, Sturrock CJ, Banda J, Torres-Martínez HH, Viana WG, et al. Moisture-responsive root-branching pathways identified in diverse maize breeding germplasm. Science. 2025:387(6734):666–673. 10.1126/science.ads5999

Schindelin J, Arganda-Carreras I, Frise E, Kaynig V, Longair M, Pietzsch T, Preibisch S, Rueden C, Saalfeld S, Schmid B, et al. Fiji: an open-source platform for biological-image analysis. Nat Methods. 2012:9(7):676–682. 10.1038/nmeth.2019

Smith LM, Bomblies K, and Weigel D. Complex Evolutionary Events at a Tandem Cluster of Arabidopsis thaliana Genes Resulting in a Single-Locus Genetic Incompatibility. PLOS Genetics. 2011:7(7):e1002164. 10.1371/journal.pgen.1002164

Song S, McDonald KJ, Bhat A, Chen MY, Morales Moreira Z, and Haney CH. FERONIA Kinase Interacting Cell Wall Sensors LRX1/2 Regulate the Plant Rhizosphere Microbiome. MPMI. 2025. 10.1094/MPMI-05-25-0064-R

Song Y, Wilson AJ, Zhang X-CC, Thoms D, Sohrabi R, Song S, Geissmann Q, Liu Y, Walgren L, He SY, et al. FERONIA restricts Pseudomonas in the rhizosphere microbiome via regulation of reactive oxygen species. Nature Plants. 2021:7(5):644–654. 10.1038/s41477-021-00914-0

Torres A, Kasturiarachi N, DuPont M, Cooper VS, Bomberger J, and Zemke A. NADH Dehydrogenases in Pseudomonas aeruginosa Growth and Virulence. Front Microbiol. 2019:10. 10.3389/fmicb.2019.00075

Tseng Y-H, Scholz SS, Fliegmann J, Krüger T, Gandhi A, Furch ACU, Kniemeyer O, Brakhage AA, and Oelmüller R. CORK1, A LRR-Malectin Receptor Kinase, Is Required for Cellooligomer-Induced Responses in Arabidopsis thaliana. Cells. 2022:11(19):2960. 10.3390/cells11192960

Wang H, Wang X, Chen J, Zhang Y, Liu J, Xia F, Li J, Wu T, and Liu L. Identification and validation of reference genes for qPCR of Pseudomonas aeruginosa L10 under varying n-hexadecane concentrations. BMC Microbiology. 2025:25(1):390. 10.1186/s12866-025-04112-2

Wang J, Ding Y, Wang J, Hillmer S, Miao Y, Lo SW, Wang X, Robinson DG, and Jiang L. EXPO, an Exocyst-Positive Organelle Distinct from Multivesicular Endosomes and Autophagosomes, Mediates Cytosol to Cell Wall Exocytosis in Arabidopsis and Tobacco Cells. The Plant Cell. 2010:22(12):4009–4030. 10.1105/tpc.110.080697

Wintermans PCA, Bakker PAHM, and Pieterse CMJ. Natural genetic variation in Arabidopsis for responsiveness to plant growth-promoting rhizobacteria. Plant Molecular Biology. 2016:90(6):623–634. 10.1007/s11103-016-0442-2

Wippel K, Tao K, Niu Y, Zgadzaj R, Kiel N, Guan R, Dahms E, Zhang P, Jensen DB, Logemann E, et al. Host preference and invasiveness of commensal bacteria in the Lotus and Arabidopsis root microbiota. Nat Microbiol. 2021:6(9):1150–1162. 10.1038/s41564-021-00941-9

Wu M and Scott AJ. Phylogenomic analysis of bacterial and archaeal sequences with AMPHORA2. Bioinformatics. 2012:28(7):1033–1034. 10.1093/bioinformatics/bts079

Yang H, Wang D, Guo L, Pan H, Yvon R, Garman S, Wu H-M, and Cheung AY. Malectin/Malectin-like domain-containing proteins: A repertoire of cell surface molecules with broad functional potential. The Cell Surface. 2021:7:100056. 10.1016/j.tcsw.2021.100056

Yeh Y-H, Panzeri D, Kadota Y, Huang Y-C, Huang P-Y, Tao C-N, Roux M, Chien H-C, Chin T-C, Chu P-W, et al. The Arabidopsis Malectin-Like/LRR-RLK IOS1 is Critical for BAK1-Dependent and BAK1-Independent Pattern-Triggered Immunity. Plant Cell. 2016:tpc.00313.2016. 10.1105/tpc.16.00313

Yuan N, Yuan S, Li Z, Zhou M, Wu P, Hu Q, Mendu V, Wang L, and Luo H. STRESS INDUCED FACTOR 2, a Leucine-Rich Repeat Kinase Regulates Basal Plant Pathogen Defense. Plant Physiol. 2018:176(4):3062–3080. 10.1104/pp.17.01266

Zamioudis C, Mastranesti P, Dhonukshe P, Blilou I, and Pieterse CMJ. Unraveling Root Developmental Programs Initiated by Beneficial Pseudomonas spp. Bacteria. Plant Physiology. 2013:162(1):304–318. 10.1104/pp.112.212597

Žárský V. Exocyst functions in plants: secretion and autophagy. FEBS Letters. 2022:596(17):2324– 2334. 10.1002/1873-3468.14430

